# Neutrophil and natural killer cell imbalances prevent muscle stem cell mediated regeneration following murine volumetric muscle loss

**DOI:** 10.1101/2021.07.02.450777

**Authors:** Jacqueline A. Larouche, Sarah J. Kurpiers, Benjamin A. Yang, Carol Davis, Paula M. Fraczek, Matthew Hall, Susan V. Brooks, Lonnie D. Shea, Carlos A. Aguilar

**Affiliations:** Dept. of Biomedical Engineering, University of Michigan, Ann Arbor, MI 48109, USA; Biointerfaces Institute, University of Michigan, Ann Arbor, MI 48109, USA; Department of Molecular & Integrative Physiology, University of Michigan, Ann Arbor, MI 48109, USA; Program in Cellular and Molecular Biology, University of Michigan, Ann Arbor, MI 48109, USA

## Abstract

Volumetric muscle loss (VML) overwhelms the innate regenerative capacity of mammalian skeletal muscle (SkM), leading to numerous disabilities and reduced quality of life. Immune cells are critical responders to muscle injury and guide tissue resident stem cell and progenitor mediated myogenic repair. However, how immune cell infiltration and inter-cellular communication networks with muscle stem cells are altered following VML and drive pathological outcomes remains underexplored. Herein, we contrast the cellular and molecular mechanisms of VML injuries that result in fibrotic degeneration or regeneration of SkM. Following degenerative VML injuries, we observe heightened infiltration of natural killer (NK) cells as well as persistence of neutrophils beyond two weeks post injury. Functional validation of NK cells revealed an antagonistic role on neutrophil accumulation in part via inducing apoptosis and CCR1 mediated chemotaxis. The persistent infiltration of neutrophils in degenerative VML injuries was found to contribute to impairments in muscle stem cell regenerative function, which was also attenuated by transforming growth factor beta 1 (*TGFβ1*). Blocking *TGFβ* signaling reduced neutrophil accumulation and fibrosis, as well as improved muscle specific force. Collectively, these results enhance our understanding of immune cell-stem cell crosstalk that drives regenerative dysfunction and provide further insight into possible avenues for fibrotic therapy exploration.

**SINGLE SENTENCE SUMMARY:** Comparison of muscle injuries resulting in regeneration or fibrosis reveals inter-cellular communication between neutrophils and natural killer cells impacts muscle stem cell mediated repair.

## BACKGROUND

The traumatic or surgical loss of a critical volume of skeletal muscle (SkM) that results in a permanent functional impairment^1^, known as volumetric muscle loss (VML), is responsible for over 90% of all muscle conditions that lead to long-term disability, and is the principle cause of nearly 10% of all medical retirements from the military^2^. VML results in reduced quality of life through pronounced disabilities^3^ ranging from reduced muscle function and atrophy to aggressive development of osteoarthritis^4^. Increasing prevalence of extremity traumas during combat since the turn of the century has incentivized increased therapeutic development for VML^5^. Towards this end, research groups have developed and tested strategies ranging from acellular biological scaffolds^6–8^ to stem cell transplants^9^, drug therapies^3,10^, and rehabilitation^11^, as well as various combinations of each^12–14^. However, each of these therapies have found limited success, with only an estimated 16% force recovery achieved^13^. These results may be due to the prolonged inflammation and inhibitive microenvironment produced by the fibrotic scar as a result of VML injury^13,15,16^. However, little information exists for why VML permanently overwhelms the regenerative capacity of SkM resulting in pathological fibrosis and degeneration.

The regenerative capacity of SkM is dependent upon a pool of resident muscle stem cells (MuSCs), also known as satellite cells^17^. Upon muscle injury, quiescent MuSCs are triggered to proliferate, differentiate into muscle progenitor cells, and fuse to form new or repair existing myofibers^17^. The immune system plays a complex role in communicating with MuSCs during the regenerative response and guides healing outcomes^18,19^. Following SkM damage, a wave of pro-inflammatory cells, including neutrophils, macrophages, and conventional T lymphocytes, infiltrate the injured site and encourage MuSC proliferation through pro-inflammatory cytokine secretion, while regulatory T cells^20,21^ balance the magnitude and duration of the inflammatory response. Once the debris has been cleared and the progenitor pool has appropriately expanded, the immune microenvironment transitions to an anti-inflammatory state to coordinate matrix rebuilding and the formation of new myotubes. As part of this transition, macrophages undergo a phenotypic switch to support myogenic progenitor differentiation and myofiber growth^22,23^. However, after VML, macrophages persist in the defect for weeks to months^13^ and their polarization switch is impaired^24^ resulting in an intermediate phenotype^25,26^. Moreover, VML triggers an increase in T helper and cytotoxic T lymphocytes^27^, which persist as long as 4 weeks post injury^26^ compared to returning to pre-injury levels within 7-10 days, as occurs in models of successful muscle regeneration^28^. Comprehensive etiological assessment after VML and how dysregulated immune signaling networks converge to influence MuSC functions remains enigmatic. Elucidating these mechanisms stands to potentiate the development of regenerative therapies.

Herein, we characterized the cellular and molecular mechanisms driving responses of SkM by comparing VML injuries that regenerate to those that degenerate and result in fibrosis. Using immunohistochemical analyses, single-cell RNA sequencing (scRNA-Seq), lineage-tracing mouse models, cellular transplants, and small molecule inhibition, we provide a resource that elucidates new cellular and molecular players post VML. We observed and validated degenerative VML injuries result in persistent infiltration of inflammatory cells, such as neutrophils, which exert lasting consequences on myogenic capacity of resident MuSCs. Further, we identified and characterized an inter-cellular communication circuit between neutrophils and cytolytic natural killer (NK) cells, which combat neutrophil accumulation via a CCL5/CCR1 axis. Inhibition of CCR1 exacerbated neutrophil accumulation in degenerative defects, while NK transplants significantly reduced neutrophil populations and enhanced healing. Finally, cell-cell communication network prediction from scRNA-Seq data suggested TGFβ-conferred MuSC impairments. Small molecule inhibition of TGFβ signaling in vivo improved tissue morphology and strength at late timepoints following degenerative VML injuries. Together, these findings enhance our understanding of cellular communication dynamics governing muscle healing outcomes.

## RESULTS

### Volumetric muscle loss induces muscle degeneration and fibrotic supplantation

To model regenerative and degenerative muscle healing outcomes, we administered bi-lateral full-thickness 2mm and 3mm punch biopsy defects to the rectus femoris of adult C57BL6/J mice^29^. Cross-sections of mouse quadriceps were stained with hematoxylin and eosin (H&E), picrosirius red (PSR), and immunohistochemically stained for laminin at 7-, 14-, 28-, and 42-days post injury (dpi) to assess muscle degeneration and fibrosis compared to uninjured controls. Consistent with previous literature^29^, the 3mm muscle defects failed to resolve by 42-dpi (subsequently termed degenerative defects), whereas regenerating myofibers filled 2mm defects (subsequently termed regenerative defects) by 28-dpi (Figure 1a). Compared to regenerative defects, degenerative defects were further characterized by higher proportions of smaller fibers (Figure 1b, n = 5-7 tissues from 3-4 mice) and significantly increased collagen deposition at 28- and 42-dpi (Figure 1c–d, Supp. Fig. 1, n = 3-7 tissues from 3-4 mice, unpaired). Combining these results shows VML defects of different sizes beget distinct regenerative trajectories.

**Figure 1.**
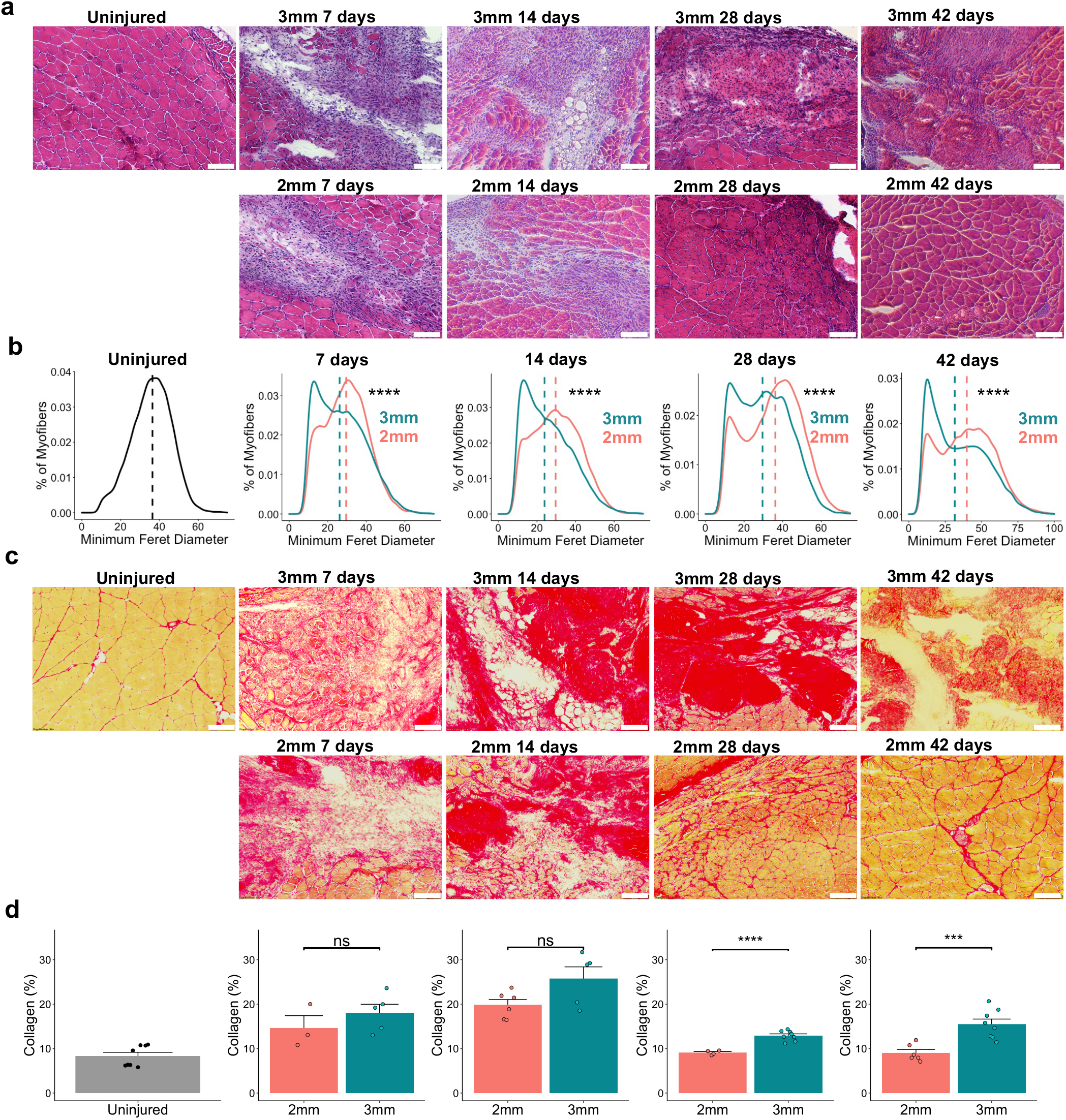
Degenerative VML defects exhibit increased fibrosis and reduced myofiber diameters. (a) Representative hematoxylin and eosin stains of uninjured quadriceps along with 2mm and 3mm defects harvested 7-, 14-, 28-, and 42-days post injury. Scale bars indicate 200 microns. Quantifications (part b) were performed on full section stitched images. Single windows of the defect area are shown for detail. (b) Portion of myofibers stratified by minimum Feret diameter shows smaller fiber size distribution following 3mm defect. ****p<0.0001 by two-sample, two-sided Kolmogorov-Smirnov test for equal distributions. D = 0.12, 0.16, 0.16, and 0.15 for 7-, 14-, 28-, and 42-dpi, respectively. Feret Diameters were calculated for 5-7 full-section images each from a distinct defect, per group, and pooled for comparison of distributions. Dashed lines indicate mean minimum Feret diameters. (c) Representative Picro Sirius Red stains of 2mm and 3mm defects throughout the time course. (d) Quantification of total collagen content showing a return to pre-injury levels following 2mm defects by 28-dpi but a significant increase and persistence following 3mm defects. Scale bars indicate 200 microns. Bars show mean ± SEM. ns denotes not significant (p>0.05). ****p < 0.0001, ***p<0.001 by two-sided t-test. n = 3-7 tissues per group. Cohen’s d = 0.7566, 1.2678, 2.2536, and 2.4264, and power = 0.1394, 0.4452, 0.9961, and 0.9894 for 7-, 14-, 28-, and 42-dpi, respectively. Quantifications were performed on full section stitched images (representatives shown in Supp. Fig. 1). Single windows of the defect area are shown for detail.

### Single cell RNA sequencing reveals alterations in cellular ecosystem in response to degenerative muscle defects

To probe the cellular and molecular drivers that contribute to the regenerative versus degenerative responses, we performed droplet-based scRNA-Seq on viable mononucleated cells isolated from uninjured quadriceps (0-dpi) as well as 7-, 14-, and 28-dpi (Figure 2a). We generated 58,405 high-quality scRNA-Seq libraries encompassing on average 1,875 genes per cell with an average read depth of 7,489 unique molecular identifiers (UMIs) per cell (Supp. Fig. 2a-b). Technical variations between batches were regressed out using Seurat v3^30^, followed by dimensionality reduction using Uniform-Manifold Approximation and Projection (UMAP)^31^ (Supp. Fig. 2c). Unsupervised Louvain clustering revealed 23 cell types (Figure 2b). Cluster-based cell-type annotation was performed by matching cluster marker genes to a muscle-specific cell taxonomy reference database using single cell Cluster-based Annotation Toolkit for Cellular Heterogeneity (scCATCH)^32^ as well as by overlaying marker gene expression with previously published reference datasets^33–35^ (Supp. Fig. 2d). We observed variation in the proportion and type of captured cells among degenerative defects compared to regenerative defects, whereby increases in neutrophils and lymphocytes were observed in degenerative defects at 7- and 14-dpi compared to cells sequenced from regenerative defects at the same time points (Figure 2c–d). Merging all time points, scRNA-Seq results suggested a 4-5-fold increase in lymphocytes and neutrophils among degenerative defects (Figure 2e). Further, substantial differences in gene expression were observed among most cell populations isolated from degenerative versus regenerative defects using the MAST toolkit^36,37^ (Figure 2f). PANTHER^38^ pathway analysis on the top 150 DEGs upregulated among degenerative defects identified increased chemokine and cytokine mediated inflammation (*Il1b, Ccl5, Ccr7, Ccr1, Rac2*, *Acta1, Actb, Actc1, Actg1, Arpc3, and Alox5ap*), T and B lymphocyte activation (*Trbc2, MHCII antigens, Cd3d, Cd79a* and *b, Ighm, and Rac2*), glycolysis (*Aldoa, Gapdh, Pgam2,* and *Bpgm*), and apoptosis signaling (*Ltb,* heat shock proteins, and *Cycs*). Most of these genes were expressed among dendritic cells (Il1b, *Ccr7, Rac2, Alox5ap*), neutrophils (*Il1b, Ltb, Ccr1, Alox5ap*), T/NK cells (*Ccl5, Ltb, Rac2, Cd3d, Trbc2,* and heat shock proteins), and B cells (*Rac2, Ccr7, Ltb, Cd79, Ighm*). We validated the proportions of T cells and NK cells using flow cytometry from regenerative and degenerative defects after 7-dpi and observed largely similar scRNA-Seq cell abundances (Supp. Fig. 3a-b, n = 6 muscles from 3 mice, unpaired). These results suggest degenerative VML defects induce stronger and sustained inflammatory responses compared to VML injuries that heal.

**Figure 2.**
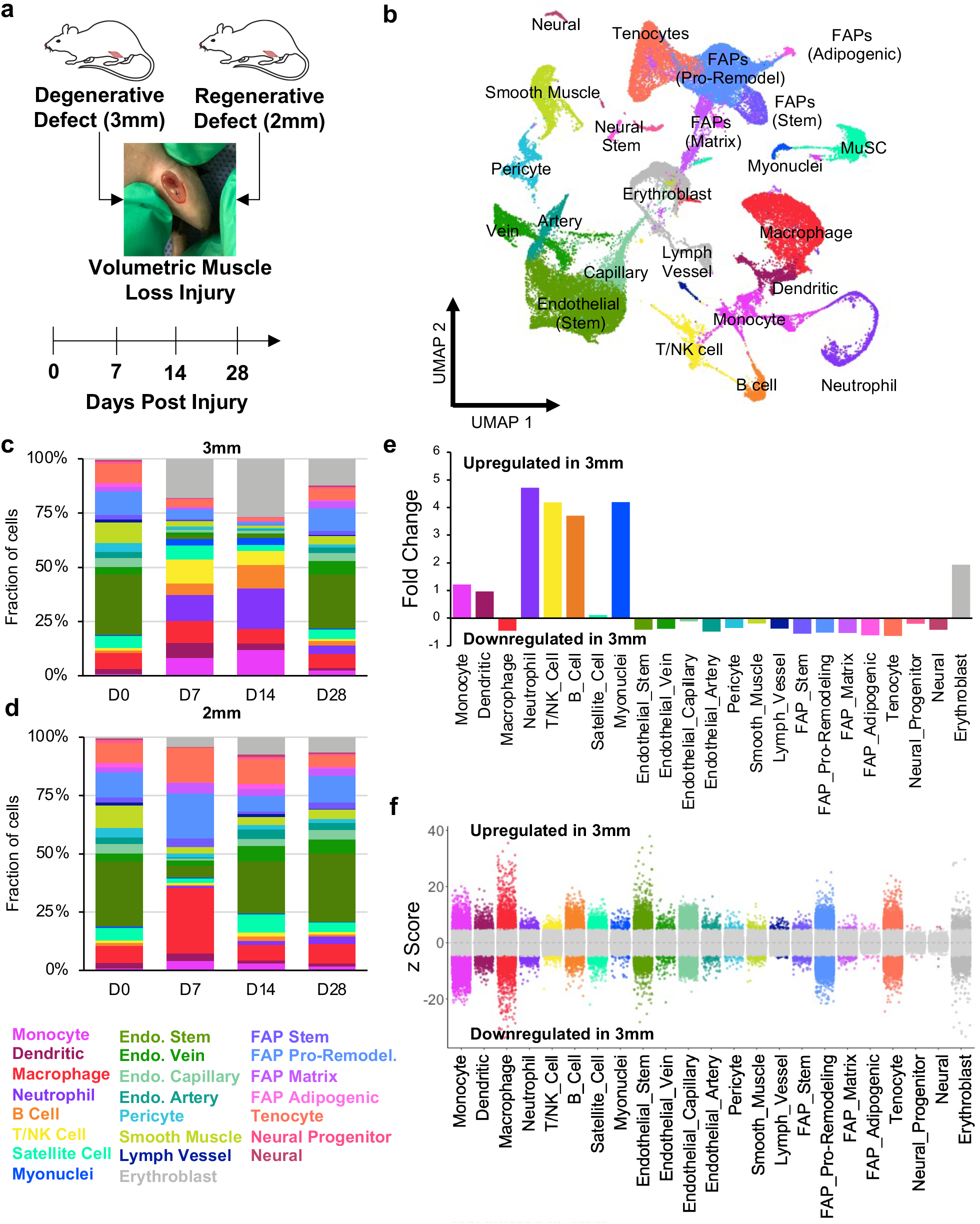
Single-Cell RNA sequencing of regenerative and degenerative muscle defects show exacerbated and persistent inflammation in injuries that do not heal. (a) Schematic of experiment, whereby adult (10-12 weeks) mice were administered 2mm or 3mm biopsy punches to their rectus femoris and humanely euthanized before injury, or 7-, 14-, or 28-dpi for scRNA-Seq analysis. (b) Dimensional reduction and unsupervised clustering of mono-nucleated cells isolated from uninjured quadriceps as well as injured quadriceps at 7-, 14-, and 28-dpi showing 23 different recovered cell types according to marker gene overlays and scCATCH cluster annotation. Two mice were pooled and sequenced for each defect size at each time-point, and between 2337 and 7500 high-quality libraries were generated for each condition. Quantification of cell abundances at each time-point sequenced following (c) 3mm and (d) 2mm VML defects shows increased and persistent inflammation among 3mm defects. (e) Fold changes in cell abundance following 3mm defects in comparison to 2mm defects merged across all timepoints shows nearly 5-fold increase in neutrophils, B cells, and T and NK cells. (f) Differential gene expression among each cell-type merged across time points and normalized to 2mm defects. Grey region indicates adjusted p-value less than 0.05. z-scores and p-values were calculated for each gene using MAST.

### Degenerative VML injuries result in variations of inflammatory cell infiltration

Our scRNA-Seq results predicted increases in neutrophils in degenerative injuries, which play a primary role in clearance of tissue debris after injury and secrete a myriad of pro-inflammatory cytokines and chemokines to recruit and activate inflammatory macrophages and T lymphocytes before being rapidly cleared^19,39^. To validate these predictions, we isolated neutrophil populations (CD11b^+^Ly6G^+^) after multiple time points following regenerative and degenerative injuries and enumerated their fractions using flow cytometry (Figure 3a). Consistent with our scRNA-Seq observations, we observed significant increases in neutrophil counts for degenerative VML injuries at 3- and 7-dpi and a trend towards increased neutrophils at 14-dpi (Figure 3b, Supp. Fig. 3c, n = 5-6 tissues from 5-6 mice, paired). To understand the mechanisms governing neutrophil accumulation following degenerative VML injuries, we employed the recently developed NicheNet^40^ algorithm. NicheNet integrates prior knowledge on ligand-target pathways with transcriptomic expression data to predict influential ligands, match them with target genes (which we defined according to differential expression across defects), and identify possible signaling mediators. NicheNet predicted that one of the top ligand-receptor interactions driving neutrophil responses in 3mm defects was CCL5/CCR1 signaling (Figure 3c). CCL5 was expressed almost exclusively in NK and T cells, while CCR1 was most highly expressed in neutrophils (Figure 3d). While over 4-fold increases in all lymphocyte sub-types were observed in degenerative defects from our scRNA-Seq data, one of the largest lymphocyte populations, and the primary source of CCL5, was NK cells (Figure 3d, Supp. Fig. 4a-b). To confirm the increased influx of NK cells into degenerative defects, we performed flow cytometry on degenerative and regenerative muscle defects at 7-dpi (Figure 3e–f, n = 6 tissues from 3 mice, unpaired) and observed about 4 times as many NK cells in degenerative defects compared with regenerative defects, which only slightly decreased after saline perfusion. Moreover, qualitative immunohistochemical stains revealed NK cells are situated at the periphery of the VML defects (Figure 3g, n = 6 tissues from 3 mice, unpaired).

**Figure 3.**
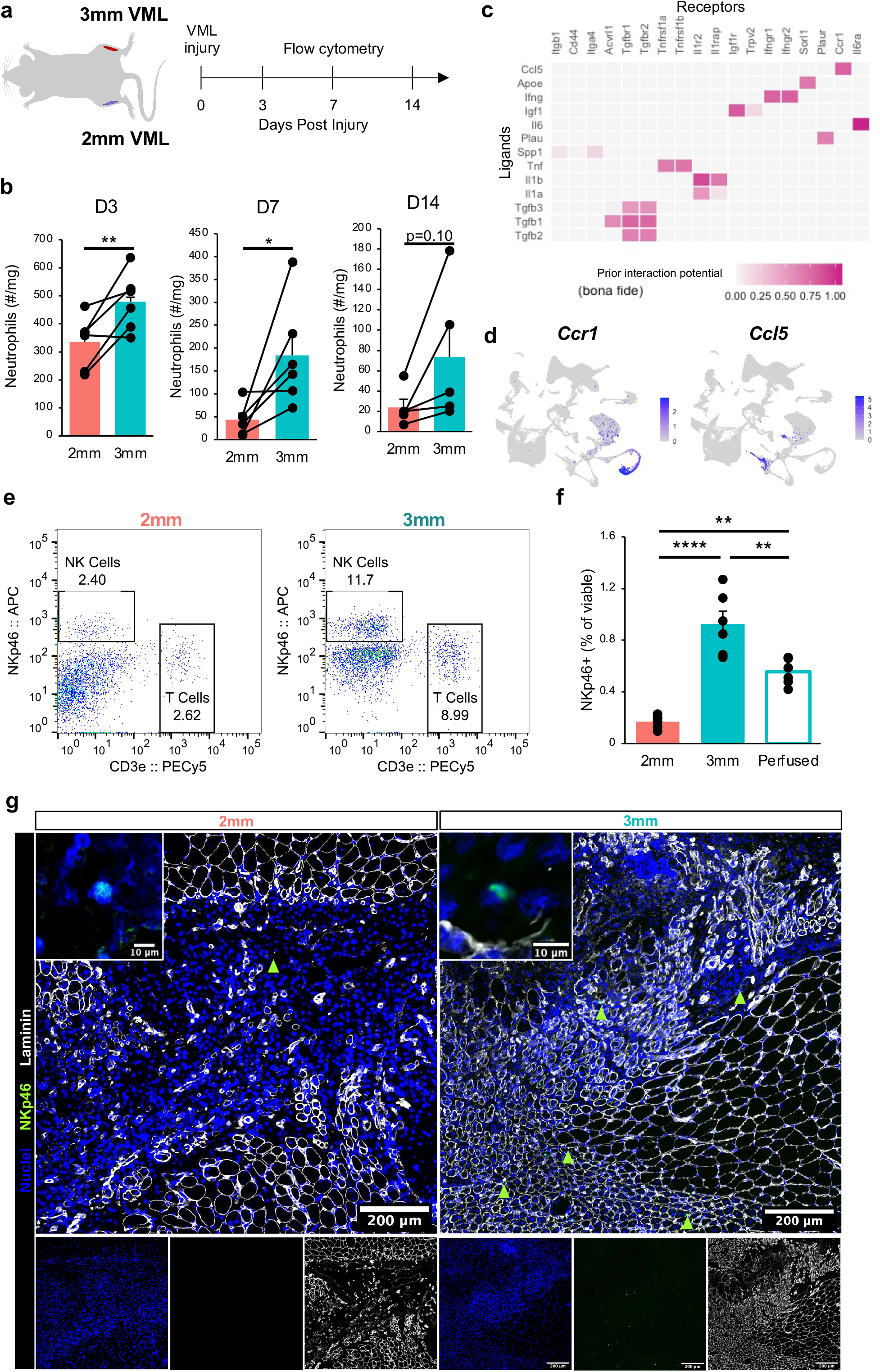
Perturbations in inflammatory cell signaling after degenerative muscle injury. (b) Time course of neutrophil infiltration in 2mm and 3mm defects quantified by flow cytometry. Mice received one 3mm and one 2mm VML defect. Boxplots show mean ± SEM. n=5-6 mice per time point. *p<0.05, **p<0.01 by two-sided, paired t-test. Cohen’s d = 2.1, 1.17, and 0.94 for 3-, 7-, and 14-dpi, respectively. (c) Top-ranked NicheNet ligand-receptor pairs based on prior literature and their prediction of downstream target gene expression. CCL5 and IFNγ are two of the top ligands predicted to influence neutrophils following 3mm defects. (d) UMAP overlays of *Ccr1* and *Ccl5* expression showing neutrophils having highest *Ccr1* mRNA expression and T/NK cells having the highest *Ccl5* mRNA expression. (e) Representative flow cytometry scatter plots showing T cell and NK cell abundance in 2mm vs 3mm VML defects at 7-dpi. (f) Flow cytometry quantification of NK cell abundance in 2mm and 3mm defects 7-dpi as well as in 3mm defects 7-dpi following whole-body saline perfusion prior to dissection. Graph shows mean ± SEM. **p < 0.001, ****p<0.0001 by one-way ANOVA and Bonferroni post-hoc analysis. n = 6 injuries per group. Effect size = 2.21. (g) Immunohistology stains qualitatively show NK cell localization primarily within the defects, and increased numbers in 3mm defects. Scale bar indicates 200um. Scale bar on inset indicates 10um.

NK cells rapidly produce and secrete pro-inflammatory cytokines^41^ and have previously been reported in SkM^21,42,43^, though their roles remain undefined. In addition to numerous NK cell marker genes (*Ncr1, Nkg7, Klre1, Klra8, Klra8*)^44^, differential gene expression analysis revealed elevated expression of pro-inflammatory molecules (*Gzma, Gzmb, Fcer1g*) and chemokines (*Ccl3, Ccl4, Ccl5*) among NK cells compared with the recovered T and NKT cells (Supp. Fig. 4c). Additionally, NK cells were observed to display strong expression of genes associated with activation and cytolytic function (*Ncr1, Klrk1, Gzma, Gzmb, Fasl*) and cytokine-secretion (*Klrc1, Ifng, Ltb*) (Supp. Fig 4d). Summing these results shows NK cells and neutrophils display enhanced infiltration following degenerative VML injuries and may communicate via a CCL5-CCR1 circuit.

### NK cells interact with neutrophils in degenerative VML injuries

To elucidate the impact of NK cells on neutrophils in VML injuries, we first co-cultured neutrophils isolated from degenerative VML defects 7-dpi with activated NK cells. Consistent with studies showing human NK cells induce contact-dependent neutrophil apoptosis^45^, we observed significantly increased neutrophil apoptosis after 4 hours (Figure 4a, Supp. Fig. 5a, n = 6 wells using neutrophils from 4 mice, unpaired). To determine whether such communication could be occurring in vivo, we next immunohistochemically evaluated co-localization of NK cells and neutrophils within VML defects at 7dpi. Over 90% of NK cells were within 50um of the nearest neutrophil, which is within previously reported maximum cytokine propagation distances up to 250 microns depending on the density of receptors^46,47^ (Figure 4b–c, n = 159 NK cells and 397 neutrophils, from 4 defects). These results suggest NK cells induce neutrophil apoptosis and are proximal to neutrophils after degenerative VML injuries.

**Figure 4.**
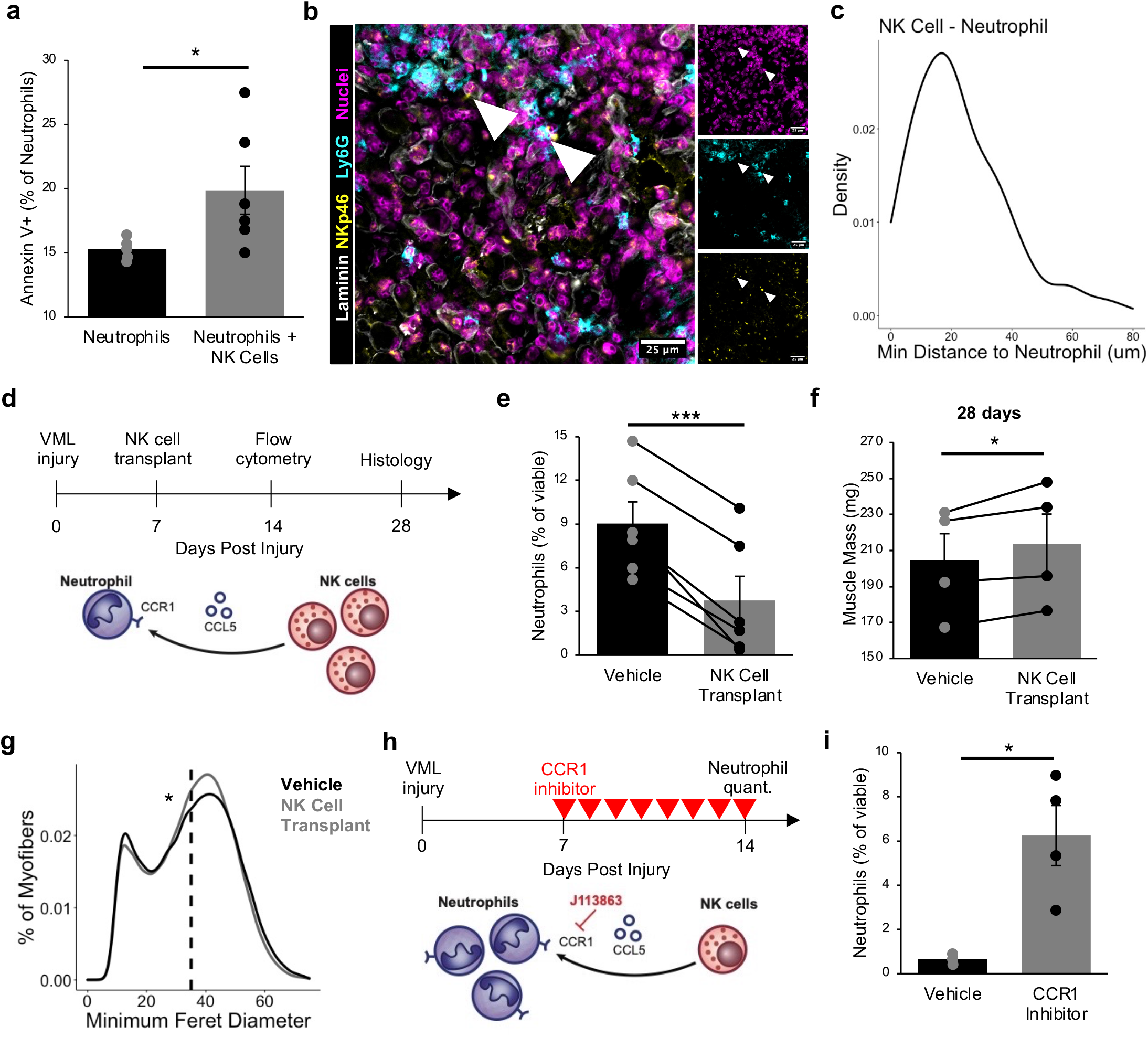
Natural killer cells interact with neutrophils in VML injuries. (a) NK cells induce neutrophil apoptosis in co-culture based on increased Annexin V+ cells, as analyzed by flow cytometry. *p<0.05. n=5-6 wells. Each well contains a pool of neutrophils FACS-enriched from four 3mm VML defects 7-dpi. NK cells were MACS-enriched from one spleen and activated in vitro for 48 hours. (b) Immunohistochemical stains for DAPI, NKp46, Ly6G, and laminin in VML defects 7-dpi show co-localization of neutrophils and NK cells. Scale bar indicates 25 microns. (c) Distributions of minimum Euclidean distances between NK cells. 92.5% of NK cells are within 50um of the nearest neutrophil. n = 159 NK cells and 397 neutrophils from 9 60X images of 4 defects. (d) Schematic whereby 150,000 in-vitro activated NK cells were transplanted into 2mm defects at 7-dpi. Contralateral control limbs received PBS injections. Neutrophil quantification and tissue morphology were assessed at 14- and 28-dpi, respectively. (e) NK cell transplant significantly reduced neutrophil abundance, quantified by flow cytometry for CD45+CD11b+Ly6G+ cells. ***p<0.001 by two-sided, paired t-test. Cohen’s d = 2.23. n = 6 muscles (N = 6 mice). Graph shows mean + standard error. (f) Quadriceps receiving an NK cell transplant had a significantly larger muscle mass, suggesting a positive impact of NK cells on regeneration. n = 4 muscles. *p < 0.05 by two-sided, paired t-test. Cohen’s d = 1.64. Graphs show mean + SEM. (g) Portion of myofibers stratified by minimum Feret diameter shows NK cell transplant reduced proportions of fibers with small minimum Feret diameters (less than 25um), which aligns with enhanced regeneration. *p<0.0001 by two-sample, two-sided Kolmogorov-Smirnov test for equal distributions. Dashed lines indicate mean minimum Feret diameters, which was approximately equal due to shift in the peak of larger fibers. n = 4 muscles. (h) Schematic of experiment design whereby a cohort of mice received 3mm defects followed by daily intraperitoneal injection of CCR1 inhibitor (J113863) between days 7 and 14 post injury. Neutrophil populations were quantified by flow cytometry (CD45+CD11b+Ly6g+) at 14-dpi. CCR1 is a receptor for CCL5 expressed on neutrophils. (i) CCR1 inhibition significantly increased neutrophil abundance in 3mm defects, to a similar extent as the NK cell transplant reduced neutrophil populations, suggesting that NK cells may locally recruit neutrophils via CCL5 secretion in order to induce contact-dependent apoptosis. *p < 0.05 by two-sided, two-sample t-test assuming equal variance. n = 3-4 mice per group, repeated twice. Cohen’s d = 2.64. Graph shows mean + SEM.

Given our results that NK cells had transcriptional profiles aligning with a cytolytic phenotype and induced neutrophil apoptosis in vitro, we reasoned that NK cell transplantation into VML defects would reduce neutrophil abundance (Figure 4d). We transplanted 150,000 *in vitro* activated NK cells into VML injured muscle at 7-dpi and quantified neutrophils and NK cells at 14-dpi using flow cytometry (Supp. Fig. 5b-c). Though the transplanted NK cells had largely cleared by 14-dpi (Supp. Fig. 5d, n = 3 tissues from 3 mice, paired), we observed a significant reduction of neutrophils in response to NK cell transplantation (Figure 4e, n = 6 tissues from 6 mice, paired). To determine if implanting NK cells impacted subsequent muscle regeneration, we histologically assessed tissues at 28-dpi. We detected increased muscle mass at 28-dpi (Figure 4f, n = 4 tissues, paired) as well as a shift towards larger myofiber minimum Feret diameters (Figure 4g, n = 4 tissues from 4 mice, paired).

To glean if NK-neutrophil communication occurs through CCR1 signaling, we administered a small molecule CCR1 inhibitor, J-113863, daily (Figure 4h). At 14-dpi, we observed a substantial increase in neutrophil abundance (Figure 4i, n = 4 tissues from 4 mice, unpaired) without significant changes to overall immune cell (CD45^+^) abundances (Supp. Fig. 5e, n = 4 tissues from 4 mice, unpaired). Taken together, these results suggest that after VML injury NK cells reduce neutrophil abundance through contact-mediated apoptosis, and that this effect is in part regulated or reinforced by CCR1 signaling.

### Degenerative VML injuries engender impairments in muscle progenitor fusion

The persistence of neutrophils after injury can have deleterious consequences for tissue repair, and failure to appropriately clear these cells from the injury site can lead to secondary damage as well as inefficient regeneration^19,39,48^. To deduce whether MuSCs may be responding to the neutrophil secretome in vivo, we sought to colocalize neutrophils with MuSCs or their progeny following VML injury. Towards this end, we administered 3mm VML defects to adult *Pax7^cre^* × *Rosa26^mTmG^* mice, which express membrane tagged green fluorescent protein (GFP) in all *Pax7*^+^ cells and their progeny following tamoxifen induction. At 14-dpi, we collected the quadriceps and immunohistochemically stained for neutrophils (Ly6G), laminin, and nuclei (Figure 5a). We observed Ly6G^+^ cells interspersed among or adjacent to GFP^+^ regenerating myofibers (Figure 5b, n = 382 neutrophils and 410 GFP^+^ fibers from 20x images of 4 defects). To understand the potential consequences of prolonged exposure of myogenic progenitors to the neutrophil secretome in the context of degenerative VML injuries, we cultured immortalized myoblasts (C2C12s) with neutrophil-conditioned growth medium for a short (1-day) or long (8-days) period, then induced differentiation and fusion (Figure 5c). We reasoned that the short exposure would mimic the quick inflammatory burst that follows regenerative defects, while the long exposure would mimic the observed significant increase in neutrophil presence beyond 7-dpi among degenerative defects. As a result of prolonged exposure to the neutrophil secretome, we observed a reduction in myoblast fusion (Figure 5d–e, n = 30 images from 3 culture wells, unpaired). Based on the observed impairments even in the absence of conditioned media during differentiation and fusion, we enriched MuSCs from degenerative and regenerative defects 28-dpi, after inflammation had largely resolved, and evaluated their ability to differentiate and fuse. We observed reductions in fusion (Figure 5f–g, n = 20 images from 5 culture wells, unpaired) among MuSCs enriched from degenerative defects compared to those enriched from regenerative defects. The negative changes in fusion were not results of differentiation impairments, as *Myog* expression was higher among MuSCs from degenerative defects compared to regenerative defects (Figure 5h, n = 3-4 culture wells, unpaired). Integrating these results demonstrates that MuSC differentiation is not impaired following VML, but fusion is partly inhibited, and that this effect may be partially conferred by extended exposure to the neutrophil secretome.

**Figure 5.**
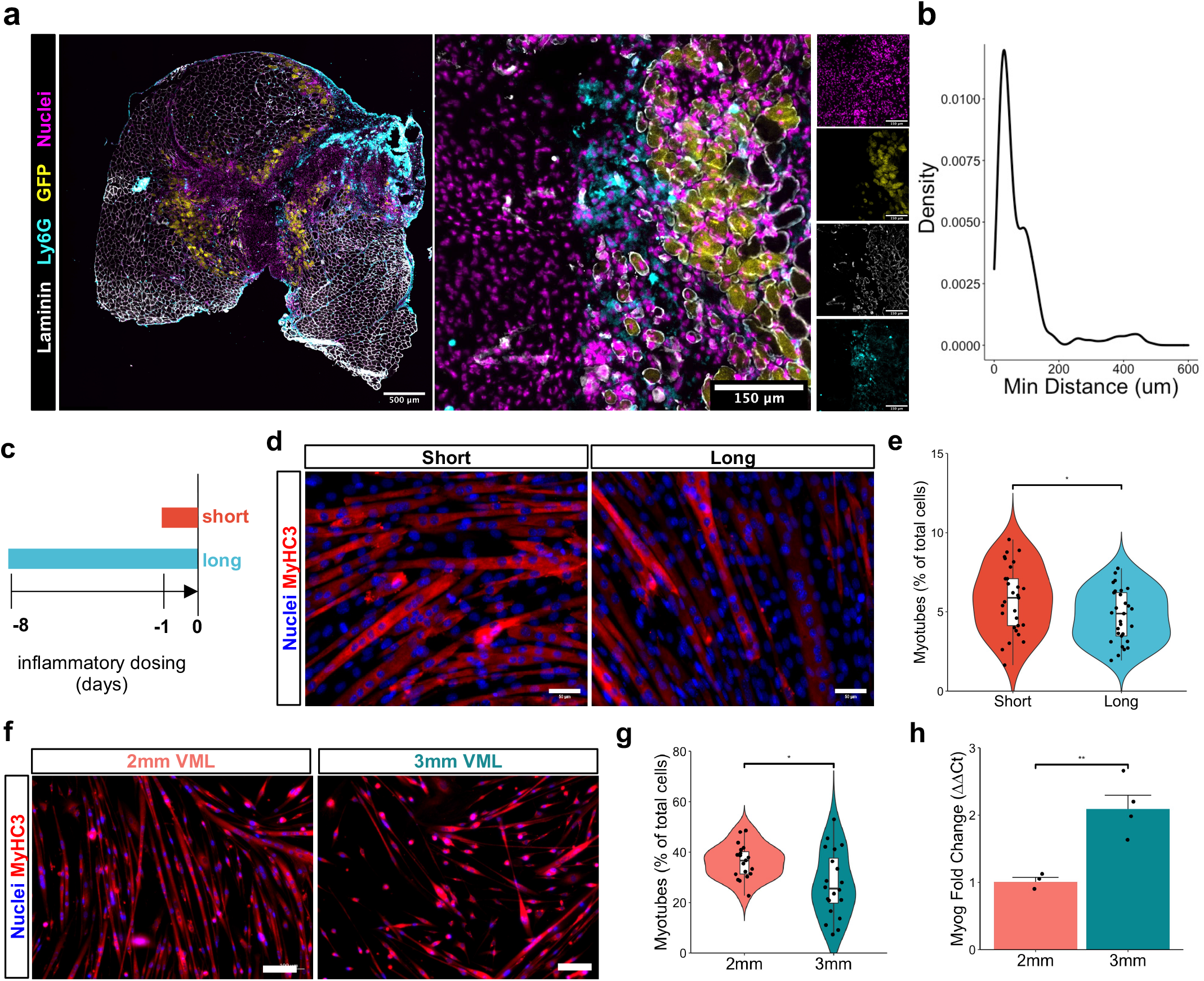
Neutrophil secretome impairs myoblast fusion in accordance with reduced muscle stem cell regenerative capacity following critical sized defects. (a) Representative image showing interspersion of neutrophils (cyan) within GFP+ regenerating myofibers (yellow) at 14-dpi. (b) Distributions of minimum Euclidean distances between neutrophils and GFP+ regenerating fibers. Over 93% of neutrophils cells are within 250um of the nearest regenerating fiber. n = 382 neutrophils and 410 GFP+ fibers from 20X images of 4 defects at 14-dpi. (c) Schematic of experiment design, whereby C2C12s are cultured in neutrophil conditioned myoblast media for 8 (long) or 1 (short) day before the addition of differentiation medium without conditioning. Fusion was assessed 72 hours later. (d) Representative immunofluorescent images of C2C12s cultured for long or short durations in neutrophil conditioned medium, then induced to differentiate for 72h in low-serum media. Blue indicates nuclei (DAPI), and red indicates embryonic myosin heavy chain (MYH3). Scale shows 50μm. (e) Quantification of the number of myotubes as a percent of total cells after short or long exposure to neutrophil conditioned medium. *p<0.05 by two-sided, two-sample t-test after removal of one outlier. n = 29 10X images taken from 3 culture wells. (f) Representative immunofluorescent images of primary myoblasts harvested 28-dpi following regenerative (2mm) and degenerative (3mm) defects and cultured in differentiation medium for 72 hours. Blue indicates nuclei (DAPI), and red indicates embryonic myosin heavy chain (MYH3). Scale bar shows 100μm (g) Quantification of fusion index according to the fraction of nuclei located in myofibers containing at least two nuclei, calculated manually. *p-value < 0.05 by two-sided, two-sample t-test, calculated from 20 10X images per defect and 3 mice per defect size, repeated twice. Cohen’s d = 0.86, power = 0.75. (h) Fold change in *Myog* expression measured by qRT-PCR following in vitro differentiation of MuSCs isolated from 2mm and 3mm defects at 28-dpi. Bars show mean ± SEM. **p<0.01 by two-sided, two-sample t-test (p = 0.02, t = 3.3897, df = 4). n=3-4. Replicates with Cq values >30 were excluded from further analysis.

To glean drivers of impairments to MuSC fusion between degenerative and regenerative defects, we analyzed differential gene expression among the MuSC scRNA-Seq cluster. The largest number of differentially expressed genes (DEGs) across all timepoints and both defects were observed among MuSCs harvested from degenerative defects at 7-dpi (455 genes, log-fold change > 1.5, adjusted p-value < 0.05). Among degenerative defects, the most upregulated genes (ordered according to adjusted p values) were associated with myogenic progenitor activation (*Itm2a, Cdkn1c, Ckb*^49^*, Atp2a1, Cebpb*^50,51^), metabolism (*Mt1* and *Mt2*^52^*, Gamt, Socs3*), fibrosis (*Sparc*)^53^, stress response, and inflammation (*Fos, Fosb, Jun, S100a8, S100a9, Prdx5, Mif* ^54^) (Supp. Fig. 6a). PANTHER^38^ pathway analysis displayed enriched terms for regulation of apoptosis (17.0% of upregulated genes), increased regulation of metabolism (35.7% of upregulated genes), and elevated an stress response (24.9% of upregulated genes). Moreover, expression of many genes associated with myoblast fusion (GO:0007520) was reduced among MuSCs in degenerative defects compared to those in regenerative defects at the same timepoint (Supp. Fig. 6b). We did not observe differences in MuSC numbers nor apoptosis at 14-dpi (Supp. Fig. 6c-d, n = 6 tissues from 6 mice, paired), however, there was a significant difference (p<0.05) in β1 integrin protein expression in degenerative defects by flow cytometry (Supp. Fig. 6e, n = 6 tissues from 6 mice, paired). Together, these results suggest the inflammatory milieu in degenerative defects induce variations in stress response, metabolism, and matrix attachment that may contribute to reductions in fusion and myogenic repair.

### Targeting predicted inter-cellular interactions alleviates muscle fibrosis by helping to resolve MuSC impairments

To elucidate which inter-cellular communication networks drive the observed differential gene expression patterns among MuSCs and impairments in fusogenic behavior, we again employed NicheNet. We observed increases in interleukin 1 (IL-1) and TGFβ1 among degenerative defects as potential drivers of the altered MuSC response (Figure 6a), influencing transcription of genes associated with attachment (*Vcam1, Itgb1*), stress response (*Fos, Fosb, Jun, C1qb, S100a8, S100a9, Hsp90aa1*), and ECM synthesis (*Col1a1, Col3a1, Fn1*) (Supp. Fig. 7a). *Il1b* was highly expressed among neutrophils and dendritic cells. *Tgfb1* was expressed in various cell types, though the most drastic upregulation across defect sizes was observed among macrophages and monocytes (Supp. Fig. 7b). TGFβ1 has previously been shown to impair MuSC fusion^55,56^ but not differentiation^57^, consistent with our in vitro observations following degenerative muscle defects (Figure 5). Moreover, TGFβ1 is one of the most potent chemokines for neutrophils^58^ and was predicted by NicheNet to be a top regulator of neutrophil populations in this system (Figure 2c). Thus, to evaluate the impact of TGFβ signaling on regeneration, we locally blocked the TGFβ signaling axis following degenerative VML injury with a TGFβ Receptor II inhibitor (ITD1) through intramuscular injection. At 28-dpi, there was a significant reduction in collagen deposition (Figure 6b–c, n = 5 tissues from 5 mice, paired). Moreover, ITD1 treatment yielded increases in maximal tetanic force, with and without normalization to muscle cross sectional area, suggesting functional improvements (Figure 6e–f, Supp. Fig. 7c-g, n = 10 tissues from 10 mice, paired). Since TGFβ1 is chemotactic for neutrophils and was among the ligands NicheNet predicted to influence neutrophils, we evaluated whether ITD1 treatment reduced neutrophil accumulation in degenerative defects using flow cytometry. In accordance with predictions, significantly fewer neutrophils were observed in ITD1 treated defects (Figure 6d, n = 4 tissues from 2 mice, unpaired). Cumulatively, our results support a network driving VML-induced muscle degeneration whereby neutrophils infiltrate and persist in degenerative defects, and with elevated TGFβ signaling, impair fusion of MuSCs^50,59^ (Figure 7).

**Figure 6.**
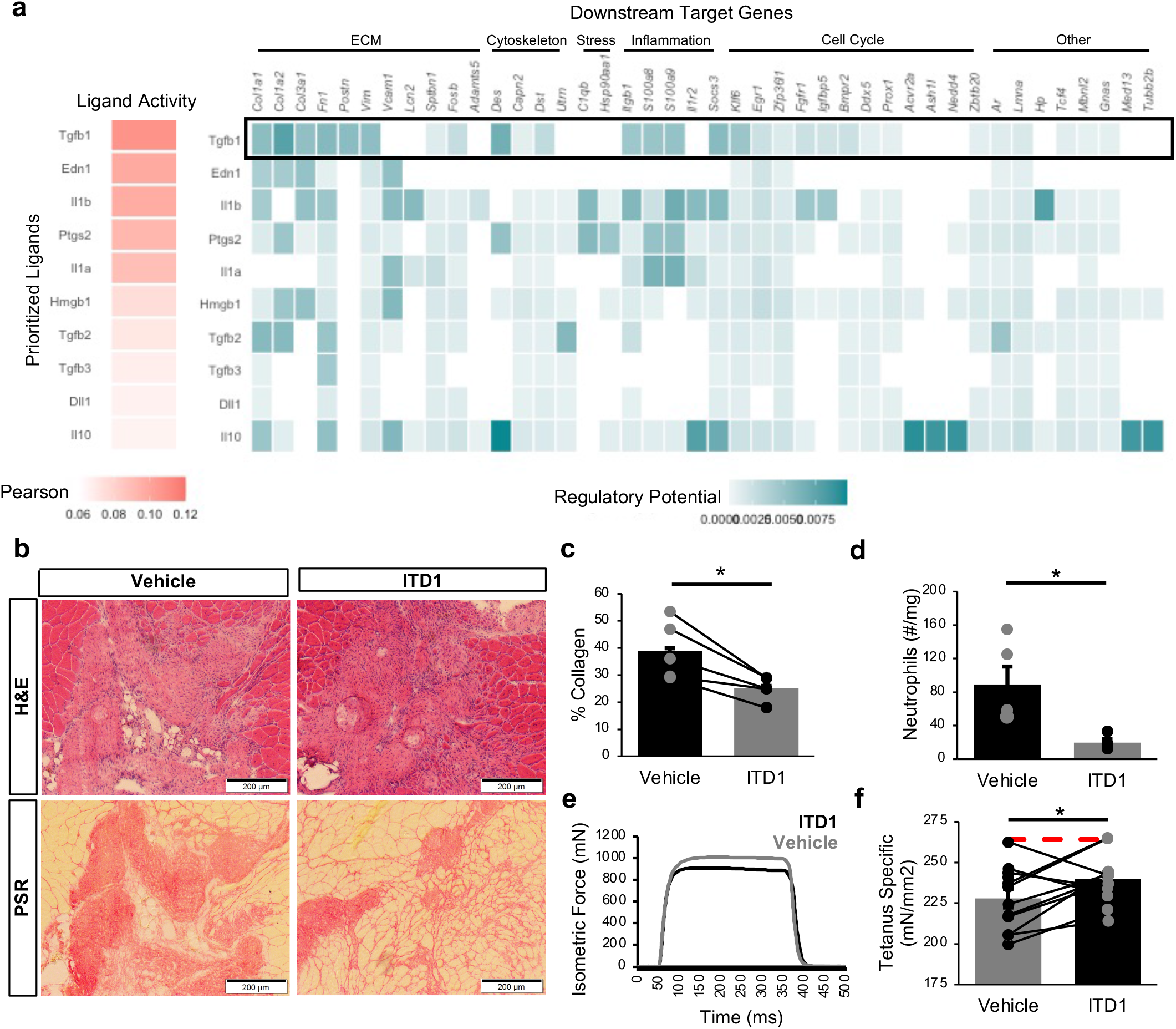
Inhibition of TGFβ signaling partially restores muscle regeneration and reduces fibrosis after VML injury. (a) NicheNet ranking of the ligands that best predict MuSC differential gene expression at 7-dpi. Pearson correlation coefficient indicates the ability of each ligand to predict the expression of differentially expressed genes. TGFβ1 is predicted to be a key contributor to MuSC dysfunction. (b) Representative H&E and PSR images from 3mm defects 28-dpi following intramuscular treatment with ITD1 (a TGFβ-signaling inhibitor) or vehicle every 3 days until 15-dpi. (c) Defects treated with ITD1 showed reduced collagen deposition. Collagen was calculated as a percent of the 10X magnification image of the defect that was stained red by PSR. n = 4 tissues per condition, where contralateral limbs were treated with vehicle. *p<0.05 by two-sided, paired t-test. Bars show mean ± SEM. Effect size = 1.83. (d) Neutrophil abundance following ITD1 treatment, as quantified by flow cytometry 14-dpi, shows TGFβ signaling inhibition reduces neutrophil recruitment. Boxplots show mean ± SEM. n=4 defects. p=0.025 by two-sided, two-sample t-test. Cohen’s d = 1.91. (e) Representative force curves for muscles treated with ITD1 or vehicle. (f) Specific force following muscle stimulation of injured tibialis anterior (TA) muscles 28-dpi is improved by ITD1 treatment. Bars show mean ± SEM, and red dashed line indicates specific force of uninjured, untreated TA muscles. p = 0.023 by two-sided, paired t-test. n = 10 muscles, Cohen’s d = 1.1.

**Figure 7.**
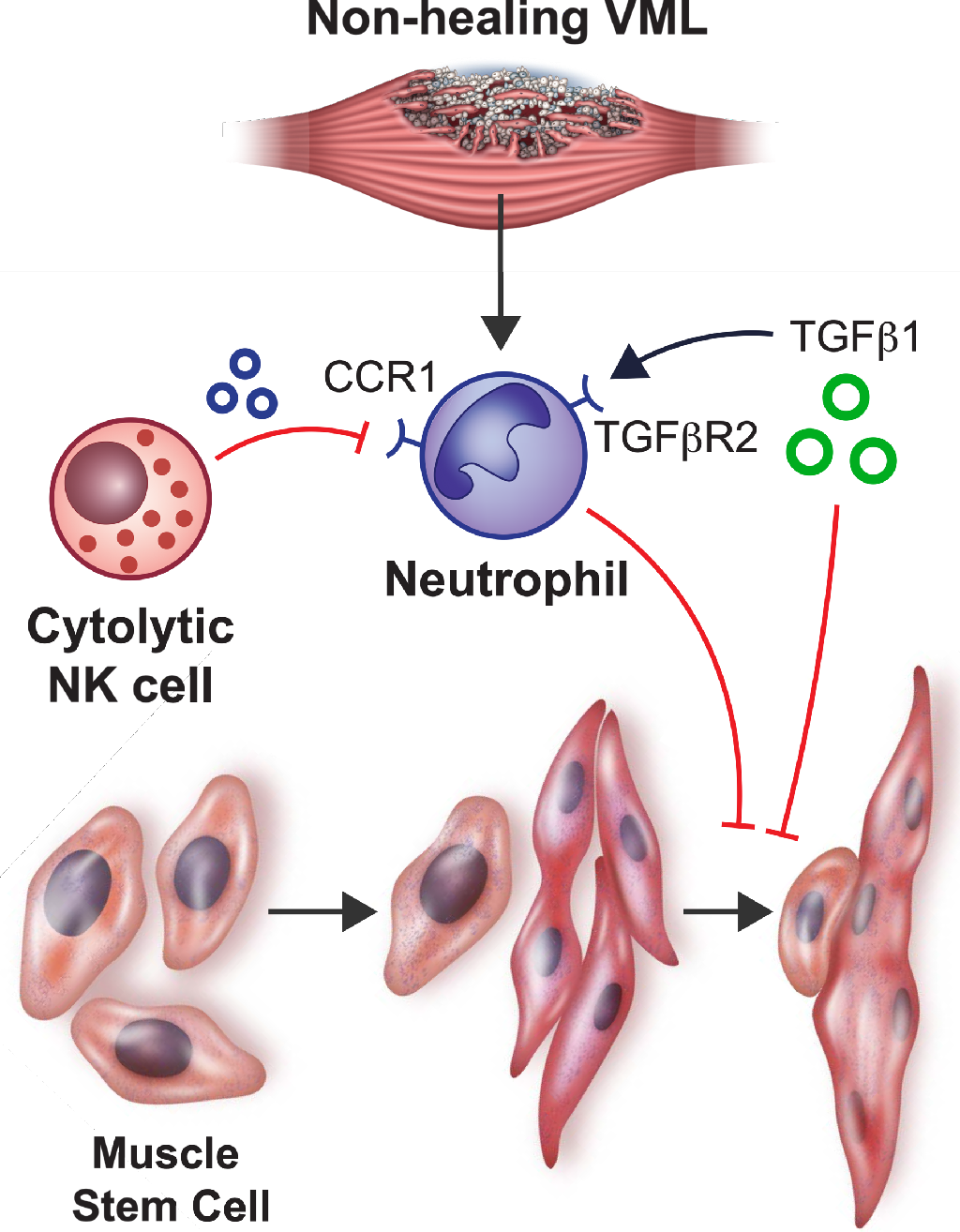
Proposed communication network following non-healing VML injuries that impairs regeneration.

## DISCUSSION

Skeletal muscle displays a remarkable ability to regenerate following injuries through coordinative actions of MuSCs^60^. Yet, the dysregulated cellular and molecular mechanisms that develop after VML and prevent MuSC-based regeneration remain elusive, limiting therapeutic efficacy and restoration of function. Understanding drivers of this pathological behavior is critical to glean the effects of new therapeutic interventions targeting these circuits as well as improve existing therapeutic modalities. Herein, we use scRNA-Seq to characterize injuries that result in degeneration and fibrosis and show that degenerative VML displays exacerbated and prolonged inflammation that negatively influences the regenerative capacity of resident MuSCs. These datasets and results offer a rich resource to further improve our understanding of the VML etiology, as well as the restorative benefits of therapies.

Regeneration of skeletal muscle after injury is critically dependent on multiple types of immune cells that remove debris, condition the injury site with inflammation to prevent infection, and signal to resident stromal and stem cells to guide repair^39^. In accordance with other muscle-regenerative injury models^39^, neutrophil abundance in regenerative VML defects peaked after injury and returned to baseline before 7 days. However, in degenerative VML injuries, neutrophils persisted beyond one week post injury and were co-located with regenerating myofibers. Collateral skeletal muscle damage caused by neutrophil-derived oxidants, as well as the ability to attenuate such damage by blocking neutrophil activity, has been extensively studied in myopathies, age-associated muscle decline, as well as various models of muscle injury and fibrosis^61^. In line with these observations^62,63^, we observed sustained exposure to neutrophil secretomes in vitro resulted in decreases in fusion of MuSCs, and correlations between reduced neutrophil abundance, larger myofiber diameters, and improved functional recovery in vivo. As such, targeting neutrophils and impacting the timeline of neutrophil clearance after VML may enhance the efficacy of regenerative therapies and alter fibrotic versus regenerative outcomes.

A corollary of increased and sustained neutrophil infiltration in degenerative VML injury was increases in a compensatory population of NK cells. We did not observe NK cells at later time points (>14 days) after degenerative injury suggesting these cells are not targeting fibrotic cells, but rather play an immunoregulatory role by targeting neutrophil populations. The enhancements in apoptosis in vitro as well as the timing of NK cell infiltration, gene expression profile and co-localization with neutrophils suggest that NK cells function to attenuate the negative inflammation induced from VML by locally recruiting and inducing neutrophil apoptosis^45,64^. This is consistent with previous reports demonstrating an anti-inflammatory role of NK cells following auto-immune myocarditis^65^. While we cannot exclude that additional factors and other cell types may also interact with NK cells^66^, crosstalk between neutrophils and NK cells has also previously been observed in cancer and chronic infections^67^, but their interactions in muscle have not been elucidated. A seminal question is why does degenerative VML injury result in sustained inflammation and fibrosis despite the presence of NK cells that act to induce neutrophil apoptosis? A clue to the mechanism of this behavior was increased TGFβ signaling in degenerative defects, which inhibits NK cell cytotoxicity^68^. Further exploration of the capacity of NK cells to control neutrophil-based inflammation in the pathological microenvironment of VML injured muscle is warranted, as well as how this crosstalk contributes to behavior of T cells and macrophages.

The role of elevated TGFβ on muscle degeneration following VML most likely results from a complex network of macrophage and fibro-adipogenic progenitor (FAP) interactions^69^. Fibro-adipogenic progenitors (FAPs) are muscle resident cells that function as important regulators of extracellular matrix (ECM) deposition. In response to exacerbated or chronic inflammation, FAPs act as pathological drivers of muscle fibrosis, intramuscular fatty infiltration, and heterotopic ossification^70^. TGFβ signaling is a critical determinant of FAP activity and response^71^, and previous efforts to reduce TGFβ after VML have shown reductions in fibrosis^71,72^. Beyond reported influences on ECM secretion by mesenchymal cells and negative impacts on MuSC activation, differentiation^3^, and fusion^56,57^, TGFβ may also be contributing to persistent neutrophil infiltration^58,73^. Moreover, it is possible that TGFβ contributes to a feed-forward degenerative loop, whereby neutrophil secretion of IL1β increases macrophage secretion of TGFβ1^74^, which then recruits more neutrophils^58,73^. This cascade has potent effects on mesenchymal stem cells and fibroblasts to synthesize collagen and TIMP1^74^, which in turn impairs MuSC regenerative capacity by manipulating activation, migration, and differentiation^75,76^. In line with this, we observed differential gene expression patterns between regenerative and degenerative MuSCs and alterations in attachment through β1 integrin^77^. These results suggest aberrant cortical tension^78^ from exacerbated ECM deposition and composition^79,80^ may alter membrane remodeling and fusogenic behavior of MuSCs^81,82^. Consistent with this and other previous findings^57^, blockade of TGFβ signaling following degenerative VML injury reduced collagen deposition and helped recover muscle strength. Because the macrophage secretome, including TGFβ1, is so critical to MuSC-mediated repair and sensitive to the local milieu^83^, additional exploration into how macrophage-MuSC crosstalk networks are dysregulated as a result of VML-induced microenvironmental changes is needed.

The cellular and molecular mechanisms driving pathological remodeling and degeneration following VML injuries remains under-examined. The approaches used in the present study begin to elucidate inter and intra-cellular signaling networks that are dysregulated as a result of critical injuries, contributing to muscle atrophy and fibrotic development. We envision these insights will provide a valuable resource for further exploration into mechanisms preventing healing after VML and therapeutics targeting those mechanisms.

## Acknowledgments

The authors thank Jesus Castor-Macias, James Markworth, and Eric Buras for advice on histological and immunohistochemical analyses, Shannon Anderson and Young Jang for assistance with the VML injury model, and the University of Michigan DNA Sequencing Core for assistance with single cell sequencing library preparation. The authors also thank other members of the Aguilar laboratory.

## Accession Code

GSE: 163376

## Author contributions

J.A.L., S.J.K., B.A.Y., C.D., P.M.F., M.H., S.V.B., and L.D.S. performed the experiments. J.A.L. analyzed the data. J.A.L., and C.A.A. designed the experiments. J.A.L. and C.A.A. wrote the manuscript with additions from other authors.

