## Supplemental Figures for "Neutrophil and natural killer cell imbalances prevent muscle stem cell mediated regeneration following murine volumetric muscle loss"

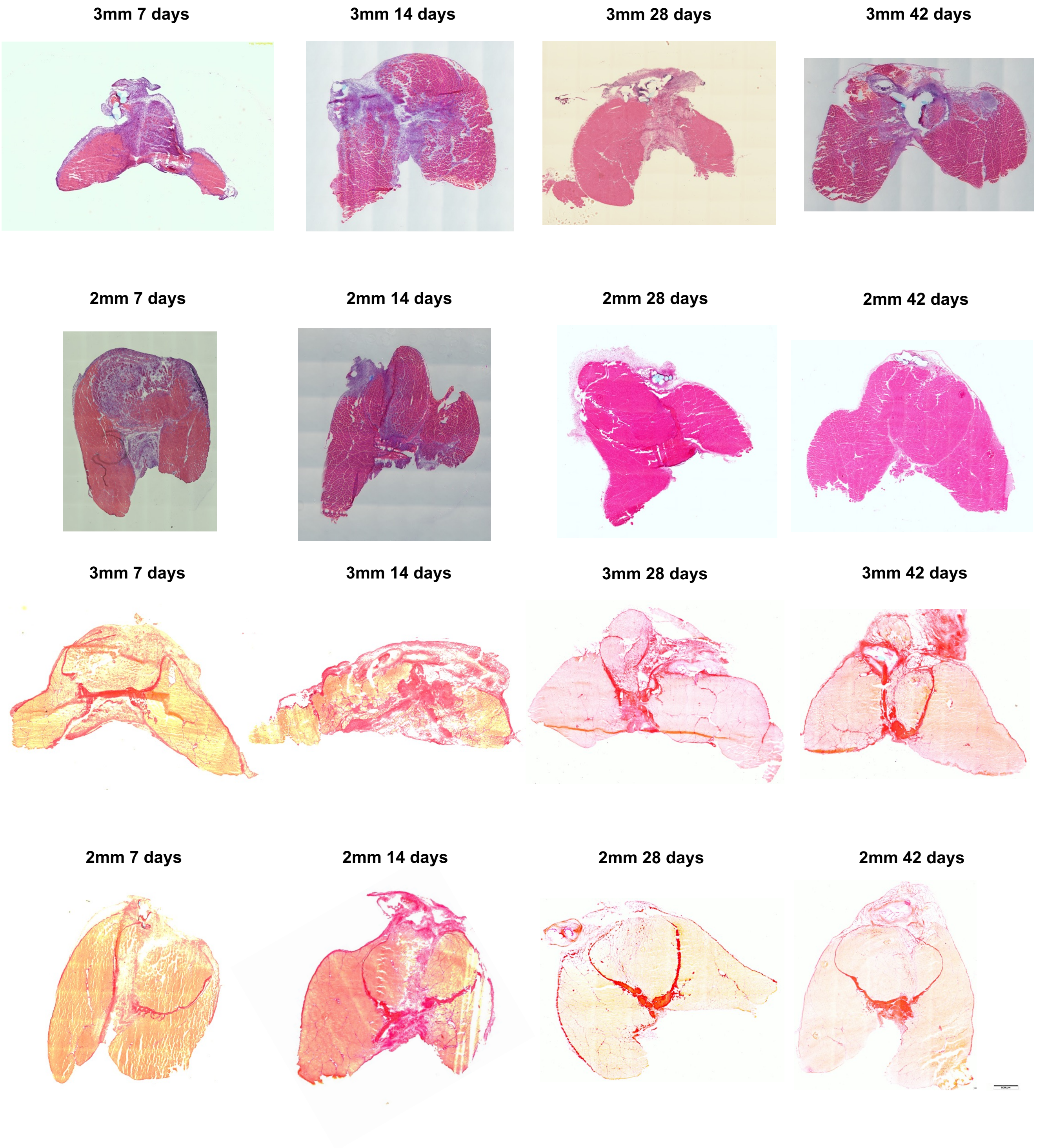


**Supplemental Figure 1.** Representative full section images of H&E (top two rows) and PSR (bottom two rows) stained tissues.

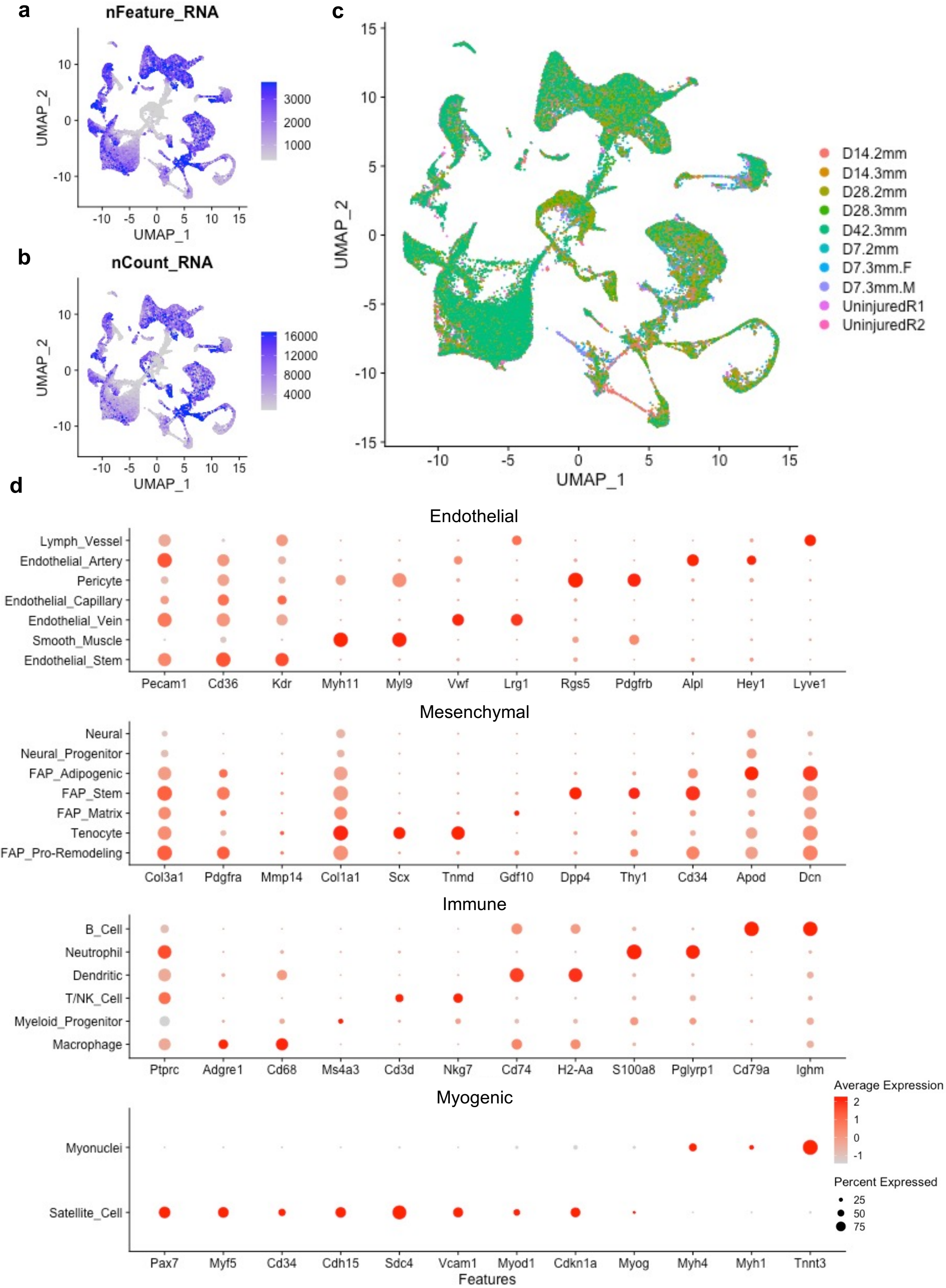


**Supplemental Figure 2. Quality control metrics for scRNA-Seq datasets and cell abundance verification.** (a) UMAP overlay of the number of genes detected per cell. (b) UMAP overlay of the total number of counts detected per cell. (c) UMAP colored by dataset following Seurat v3 batch correction. (d) Dot plots of marker gene expression showing different clusters of cells used for annotation.

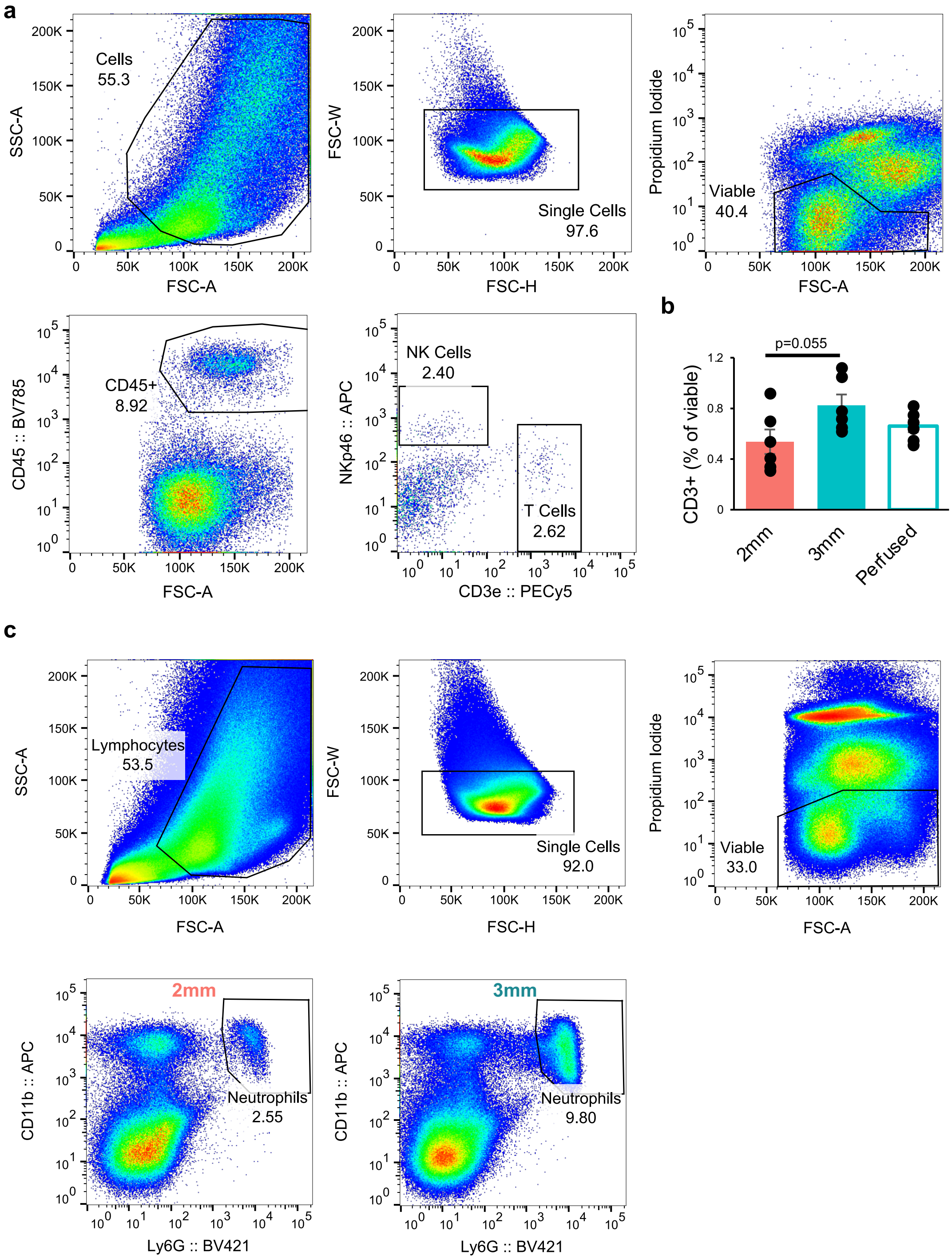


**Supplemental Figure 3. Flow cytometry verification of cell abundance.** (a) Representative flow cytometry scatterplots showing gating strategy for T cells and NK cells. (b) Flow cytometry quantification of CD3+ T cell abundance at 7-dpi. Bars show mean ± SEM. No significant difference was observed by one-way ANOVA between 2mm and 3mm defects, nor after 3mm defects with PBS perfusion immediately prior to muscle harvest, though there is a trend towards more t cells in degenerative defects. n = 6 muscles per group. NK cell quantifications by flow cytometry are shown in Figure 2e. (c) Representative flow cytometry scatterplots showing gating strategy for neutrophils. Neutrophil abundance quantifications by flow cytometry are shown in Figure 2a.

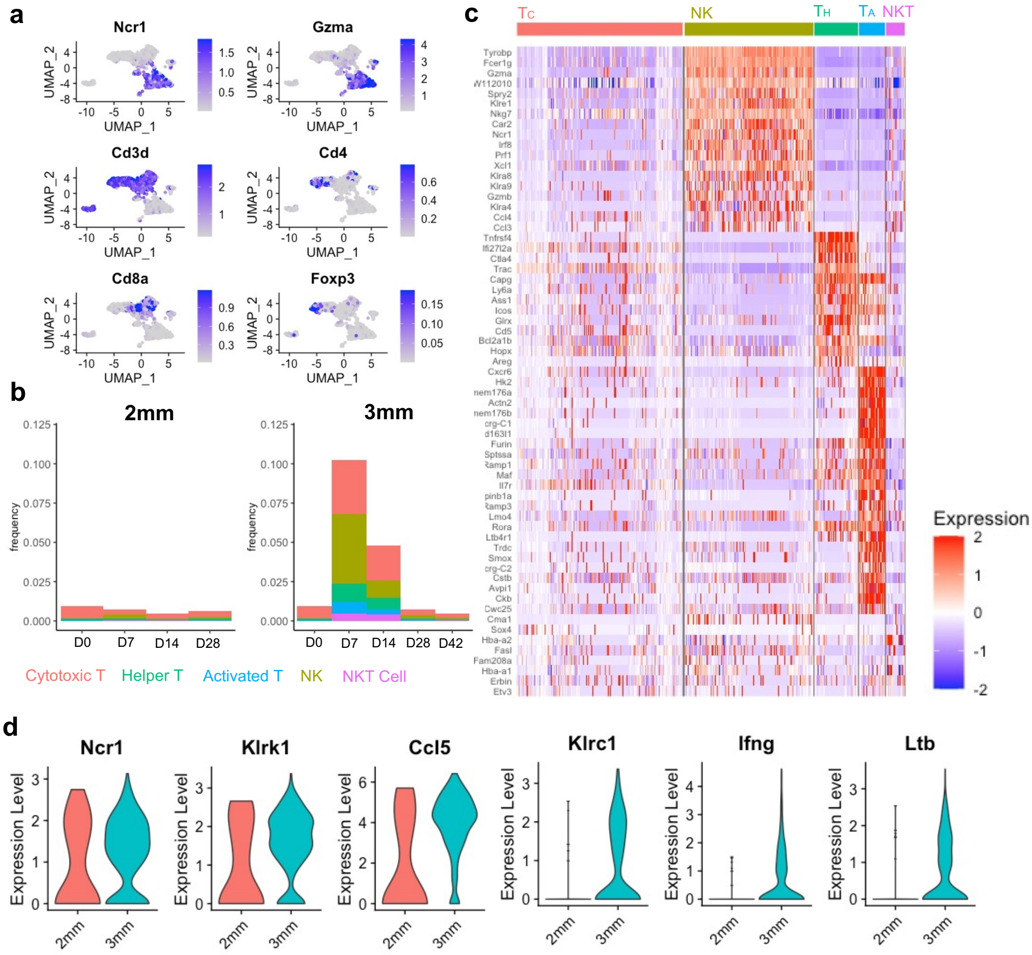


**Supplemental Figure 4. Identification and characterization of defect-infiltrating NK cells.** (a) Expression of canonical T and NK cell marker genes overlayed onto UMAP dimensionally reduced NK/T cell cluster shows the presence of NK cells, NKT cells, and three subsets of T cells (T helper cells, cytotoxic T cells, and activated T cells). (b) Lymphocyte subtypes as a fraction of total cell population at each time point post injury in 2mm vs 3mm defects from scRNA-Seq. (c) Heatmap of differentially expressed genes across lymphocyte populations in VML defects, grouped by cell type. Tc = cytotoxic (CD8+) T cell, Th = helper (CD4+) T cell, Ta = activated T cell, NK = natural killer cell, NKT = natural killer T cell. (d) Expression of *Ccl5*, cytolytic- (*Ncr1, Klrk1*) and cytokine secretion- (*Klrc1, Ifng, Ltb*) associated NK cell markers suggests in NK cells isolated from 2mm versus 3mm defects.

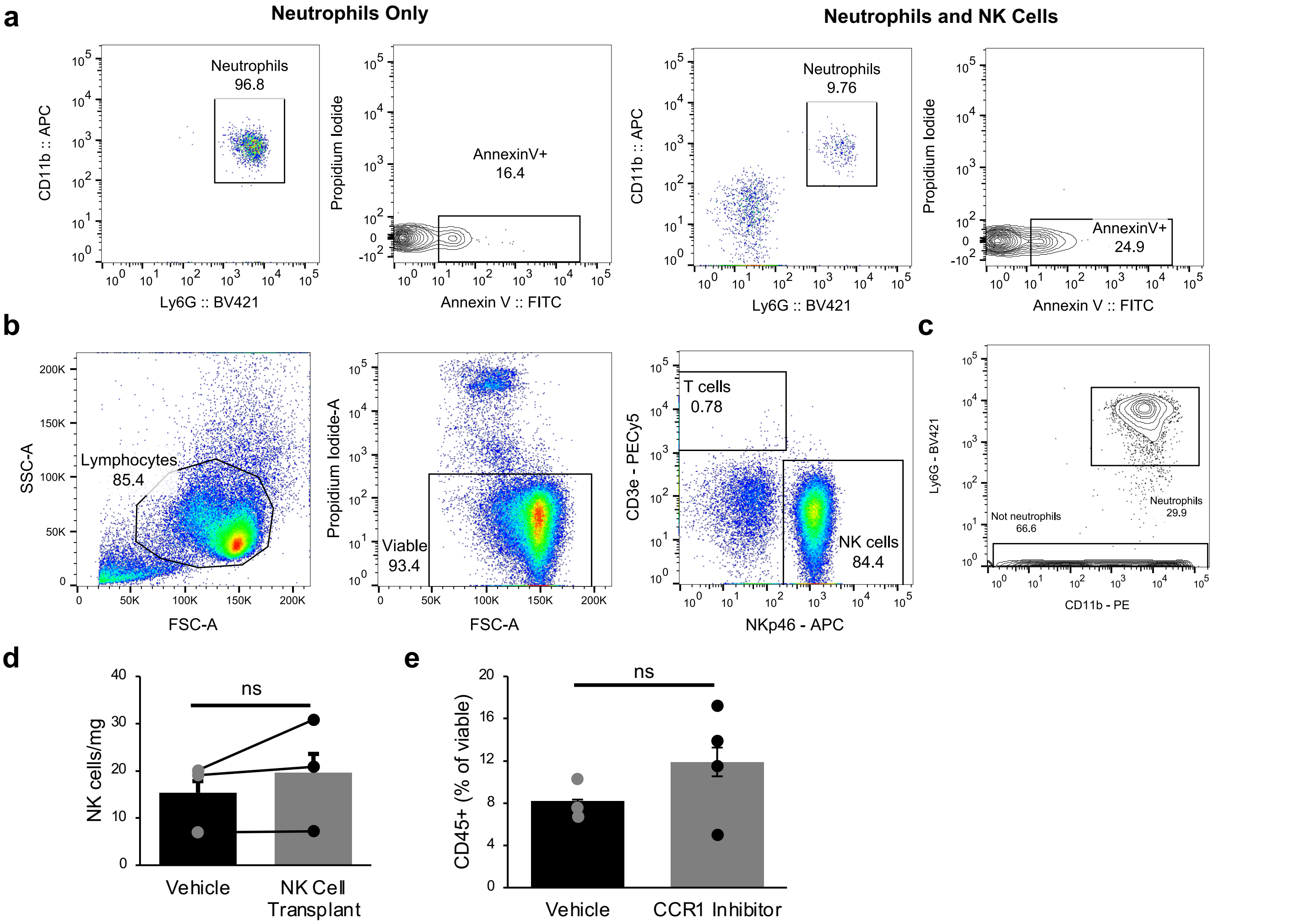


**Supplemental Figure 5. Isolation and Transplantation of Natural Killer Cells.** Reference to figure 4. (a) Representative flow cytometry scatter plots showing gating for Annexin V co-culture staining. (b) Flow cytometry shows magnetically enriched NK cell population is ~85% pure based on NKp46 expression and 93% viable. (c) Flow cytometry gating for neutrophils and lymphocytes 14-dpi and 7 days post NK cell transplant. Cells were previously gated on PI exclusion and CD45 expression. (d) Transplanted NK cells had cleared from the muscle by 14-dpi (7 days post transplant). Graph shows mean ± SEM. ns denotes not significant ($p = 0.8059$) by two-sided, paired t-test. n = 3 muscles per group. (e) CCR1 inhibition resulted in a trend towards increased immune cells at 14dpi, though the 3.5 percent increase in means (8.21% in vehicle, 11.9% in CCR1i) is smaller than the increase in neutrophils as a percent of viable (6% increase), suggesting CCR1i does not largely impact abundance of other immune cell types. $p > 0.05$ by two-sided, two-sample t-test assuming equal variance. n = 3-4 mice per group, repeated twice. Cohen's d = 0.89. Graph shows mean + SEM.

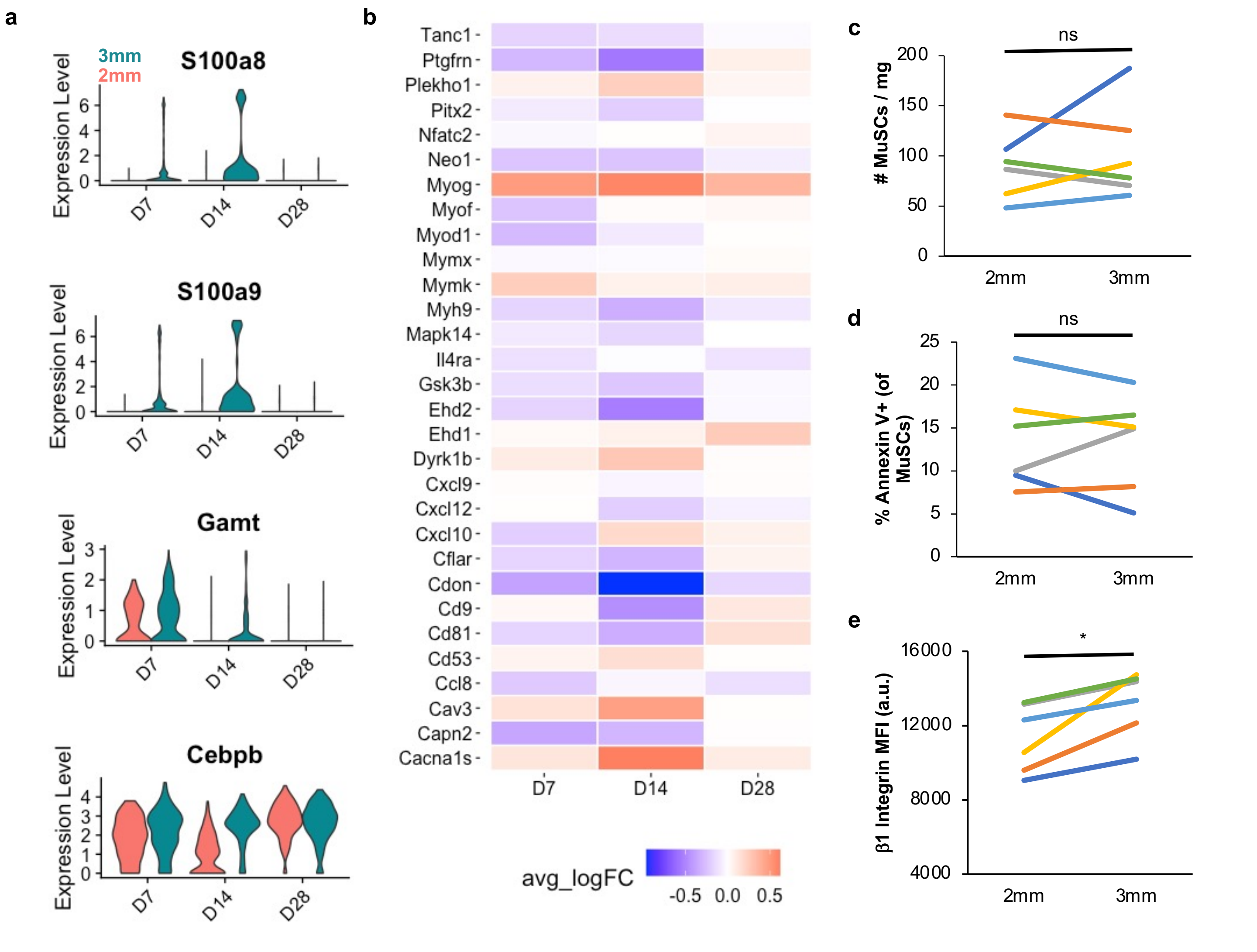


**Supplemental Figure 6. Comparison of MuSCs apoptosis and matrix attachment.** (a) Differentially expressed genes across each condition among the MuSC scRNA-Seq cluster. (b) Heatmap of genes associated with myoblast fusion among MuSCs at different timepoints. Average fold change of each gene was calculated for MuSCs from 3mm defects in relation to MuSCs from 2mm defects at each timepoint, such that red bars indicate increased expression among MuSCs from 3mm defects and blue bars indicate reduced expression at that timepoint. (c) Number of MuSCs in quadriceps 14-dpi, quantified by flow cytometry ($CD45^-$ $Ter119^-$ $CD31^-$ $Sca1^-$ $CD11b^-$ $PI^-$ $CXCR4^+$ $B1int^+$). n = 6 muscles per group, each mouse receiving one 2mm defect and one 3mm defect. No significant difference across defect sizes by paired t-test (p = 0.45). Each line represents one mouse. (d) Percent of viable ($PI^-$) MuSCs that are apoptotic (Annexin $V^+$) 14-dpi, as determined by flow cytometry. n = 6 muscles per group, each mouse receiving one 2mm defect and one 3mm defect. No significant difference across defect sizes by paired t-test (p = 0.78). Each line represents one mouse. (e) Integrin β1 expression is lower in MuSCs harvested from regenerative defects 14-dpi based on mean fluorescent intensity (MFI). n = 6 muscles per group. Significant difference across defect sizes by paired t-test (p = 0.01). Each line represents one mouse.

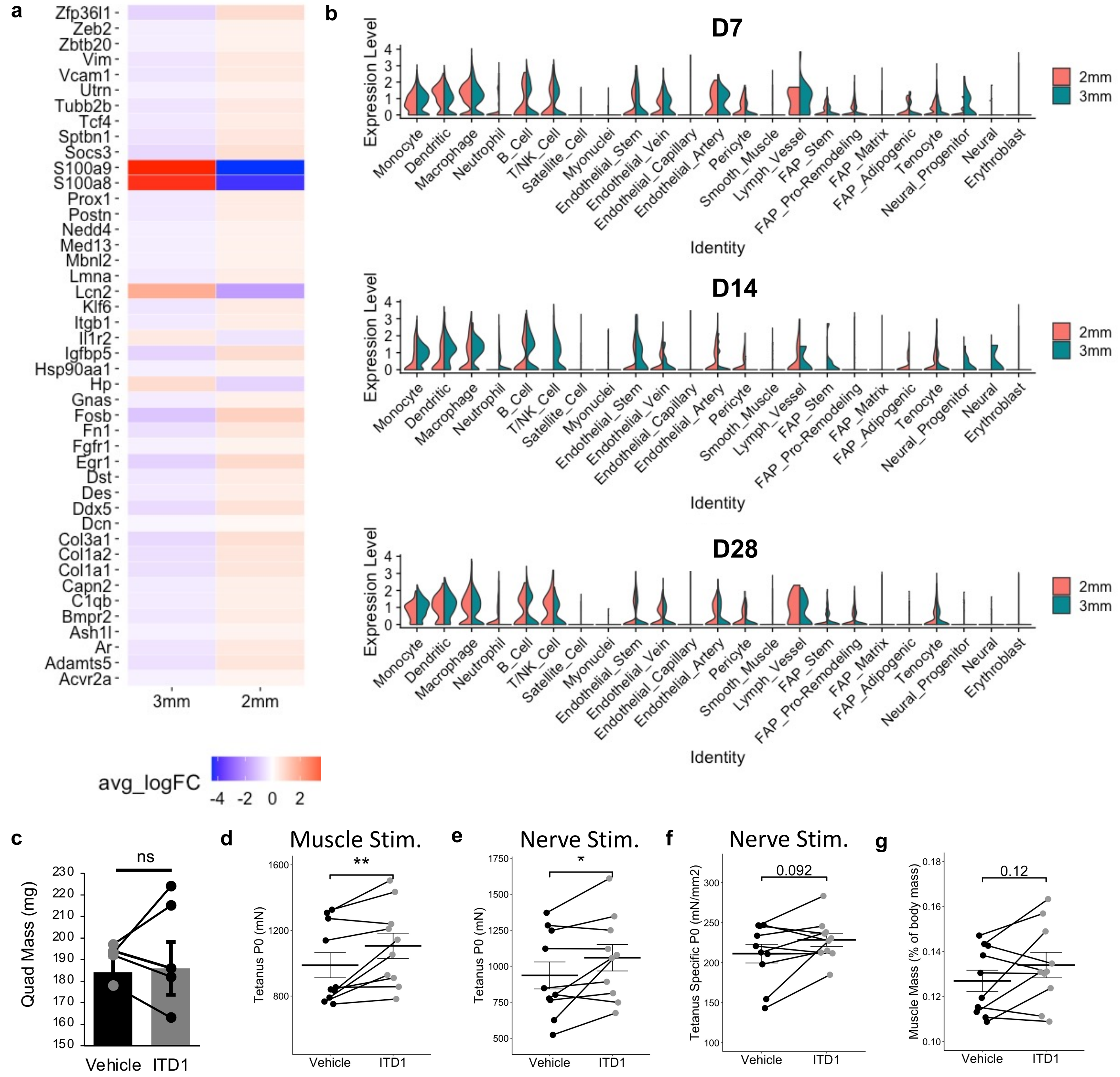


**Supplemental Figure 7.** (a) Average log fold change of gene expression among NicheNet predicted target genes at 7-dpi. (b) Average TGFβ1 expression among each cell type across the time-course shows drastic upregulation primarily among macrophages and monocytes in 3mm defects. (c) ITD1 treatment had no effect on quadricep mass at 28-dpi. (d) Impact of ITD1 on total force when the muscle was stimulated. **p<0.01 by two-sided, paired t-test. n=10 muscles (6 males, 4 females). (e) ITD1 treatment significantly impacted maximum tetanic force when the nerve was stimulated, (f) but it did not significantly impact specific force. *p<0.05 by two-sided, paired t-test. n=10 muscles (6 males, 4 females). (g) ITD1 treatment had a mild, but not significant, impact on TA muscle mass at 28-dpi. p=0.12 by two-sided, paired t-test. n=12 muscles (6 males, 6 females).
