## Supplementary material for "Neutrophil and natural killer cell imbalances prevent muscle stem cell mediated regeneration following murine volumetric muscle loss": Materials and Methods

KEY RESOURCES TABLE

| REAGENT or RESOURCE | SOURCE | IDENTIFIER |
|---|---|---|
| **Antibodies** | | |
| Rat anti-mouse Ly6G | Abcam | ab25377; RRID:AB_470492 |
| APC anti-mouse/human CD11b | BioLegend | 101212; RRID:AB_312795 |
| PE/Cy5 anti-mouse CD3e | BioLegend | 100310; RRID:AB_312675 |
| AF488 anti-mouse CD45 | BioLegend | 103149; RRID:AB_2564590 |
| FITC anti-mouse CD68 | BioLegend | 137006; RRID:AB_10578412 |
| APC anti-mouse CD335 (NKp46) | BioLegend | 137608; RRID:AB_10612758 |
| BV421 anti-mouse Ly-6G | BioLegend | 127627; RRID:AB_10897944 |
| APC anti-mouse Ly-6A/E (Sca-1) | BioLegend | 108112; RRID:AB_313349 |
| APC anti-mouse CD45 | BioLegend | 103112; RRID:AB_312977 |
| APC anti-mouse TER-119 | BioLegend | 116212; RRID:AB_313713 |
| PE anti-mouse/rat CD29 | BioLegend | 102208; RRID:AB_312885 |
| Biotin Rat Anti-Mouse CD184 | BD Bioscience | 551968; RRID:AB_394307 |
| Streptavidin PE-Cyanine7 | Thermo Fisher | 25-4317-82; RRID:AB_10116480 |
| Goat anti-mouse NKp46/NCR1 | R&D Systems | AF2225; RRID:AB_355192 |
| Rabbit anti-mouse laminin 1+2 | Abcam | ab7463; RRID:AB_305933 |
| GFP monoclonal antibody [GF28R] | Invitrogen | MA5-15256; RRID:AB_10979281 |
| Donkey anti-goat IgG (H+L), Alexa Fluor 488 | Abcam | ab150129; RRID:AB_2687506 |
| Goat anti-rabbit IgG (H+L), Alexa Fluor 647 conjugate | Thermo Fisher | A27040; RRID:AB_2536101 |
| Goat anti-rat (H+L), Alexa Fluor 647 conjugate | Thermo Fisher | A21247; RRID:AB_141778 |
| Goat anti-mouse (H+L), Alexa Fluor 488 conjugate | Thermo Fisher | A28175; RRID:AB_2536161 |
| Goat anti-rabbit (H+L), Alexa Fluor 555 conjugate | Thermo Fisher | A21247; RRID:AB_141778 |
| Goat anti-mouse H+L, Alexa Fluor 555 conjugate | Thermo Fisher | A28180; RRID:AB_2536164 |
| **Chemicals, Peptides, and Recombinant Proteins** | | |
| Dispase II (activity ≥0.5 units/mg solid) | Sigma | D4693-1G |
| Collagenase Type II (654 U/mg, non-specific proteolytic activity 487 U/mg) | Life Technologies | 17101015 |
| DMEM, high glucose, pyruvate | Life Technologies | 11995065 |
| DMEM, high glucose, pyruvate, GlutaMAX | Life Technologies | 10569010 |
| Ham's F-10 Nutrient Mix | Life Technologies | 11550043 |
| RPMI 1640 | Gibco | 11875119 |
| Fetal Bovine Serum | Life Technologies | 10437028 |
| Normal Goat Serum | Abcam | Ab7481; RRID:AB_2716553 |
| HBSS, no calcium, no magnesium, no phenol red | Life Technologies | 14175145 |
| Propidium Iodide - 1.0 mg/mL Solution in Water | Life Technologies | P3566 |
| Mouse on Mouse blocking reagent | Vector Labs | MKB-2213 |
| Tissue Plus O.C.T Compound | Fisher Scientific | 23-730-571 |
| Hematoxylin | Ricca Chemical Company | 3530-16 |
| Eosin | EMD-Millipore | 588X-75 |
| Magnesium Sulfate Heptahydrate | Sigma Aldrich | 63138-250G |
| Sodium Bicarbonate | Sigma Aldrich | S5761 |
| SafeClear II | Fisher Scientific | 23-044192 |
| Direct Red 80 | Fisher Scientific | AAB2169306 |
| Picric Acid | Sigma Aldrich | P6744-1GA |
| Glacial Acetic Acid | Sigma Aldrich | BP2401-500 |
| Xylenes | Sigma Aldrich | 534056-4L |
| Permount | Fisher Scientific | SP15-100 |

| | | |
|---|---|---|
| Bovine Serum Albumin | Fisher Scientific | BP9703-100 |
| 0.5M EDTA | Invitrogen | 15575-038 |
| Sodium Azide | Sigma Aldrich | 71289 |
| 4% paraformaldehyde in PBS | Thermo Fisher | J19943-K2 |
| Matrigel | BD Biosciences | 356234 |
| Horse Serum | Gibco-Invitrogen | 26050088 |
| Fibroblast Growth Factor basic | Gibco-Invitrogen | PHG0263 |
| Prolong Diamond | Thermo Fisher | P36965 |
| 0.25% Trypsin EDTA | Gibco-Invitrogen | 25200072 |
| Penicillin Streptomycin | Gibco-Invitrogen | 15640055 |
| QIAzol | Qiagen | 79306 |
| Tween-20 | Sigma Aldrich | P1379 |
| TritonX-100 | Sigma Aldrich | T8787 |
| SYBR Green PCR MasterMix | Thermo Fisher | 4309155 |
| PrimeTime Mouse GAPDH Primer | Integrated DNA Technologies | Mm.PT.39a1 |
| PrimeTime Mouse Myog Primer | Integrated DNA Technologies | Mm.PT.58.6732917 |
| Hoechst 33342 | Thermo Fisher | H3570 |
| J-113863 | Tocris Biosciences | 2595 |
| Cremophor EL | Sigma Aldrich | 238470 |
| Recombinant Mouse IL-15R alpha Fc Chimera Protein, CF | R&D Systems | 551-MR-100 |
| Recombinant murine IL-15 | Peprotech | 210-15 |
| RPMI | Invitrogen | 61870036 |
| FITC Annexin V | BioLegend | 640906 |
| Annexin V Binding Buffer | BioLegend | 422201 |
| ITD-1 | R&D Systems | 5068/10 |
| DMSO | Fisher Scientific | D128-4 |
| LS Columns | Miltenyi | 130-042-401 |
| Liberase TL | Sigma Aldrich | 5401020001 |
| UltraComp eBeads | Fisher Scientific | 01-2222-42 |
| **Critical Commercial Assays** | | |
| Satellite Cell Isolation Kit, mouse | Miltenyi | 130-104-268 |
| NK Cell Isolation Kit, mouse | Miltenyi | 130-115-818 |
| Neutrophil Isolation Kit, mouse | Miltenyi | 130-097-658 |
| SuperScript III First-Strand Synthesis Kit | Thermo Fisher | 18080051 |
| Qiagen miRNeasy Micro Kit | Qiagen | 217084 |
| Qubit RNA HS Assay | Thermo Fisher | Q32852 |
| Single cell 3’ Library & Gel Bead Kit v2 and v3 | 10x Genomics | 120267 |
| **Deposited Data** | | |
| scRNA-Seq datasets | This Manuscript | GSE 163376 |
| **Experimental Models: Organisms/Strains** | | |
| C57BL/6J wild-type mice (2-3 months) | Jackson Labs | Stock 000664 |
| Pax7Cre$^{ER/+}$;Rosa26$^{mTmG/+}$ mice (3 months) | University of Michigan | Jackson stock 017763 crossed with stock 007676 |

| | | |
|---|---|---|
| Pax7Cre$^{ER/+}$;Rosa26$^{dTomato/+}$ fmice (3 months) | University of Michigan | Jackson stock 017763 crossed with stock 007676 |
| **Software and Algorithms** | | |
| CellRanger v3.0.0 | 10x Genomics | https://support.10xgenomics.com/single-cell-gene-expression/software/downloads |
| R v4.0.2 | The R Foundation for Statistical Computing | https://www.r-project.org/ |
| Jupyter v4.4.0, notebook v5.7.8 | Project Jupyter 2019 | https://jupyter.org/ |
| Python v3.6.5 | Python Software Foundation | https://www.python.org |
| MATLAB_R2019b | MathWorks | https://www.mathworks.com/products/matlab.html |
| Seurat v3.1.2 | Butler et al. 2019 | https://satijalab.org/seurat/ |
| SeuratWrappers v0.3.0 | | https://github.com/satijalab/seurat-wrappers |
| scCATCH v2.0 | Shao 2020 | https://github.com/ZJUFanLab/scCATCH |
| NicheNet v1.0.0 | Bonnardel et al. 2020 | https://github.com/saeyslab/nichenetr |
| Cowplot v1.0.0 | | |
| ggplot2 v3.2.1 | Wickham 2016 | https://ggplot2.tidyverse.org |
| dplyr v0.8.3, v1.0.2 | Wickham 2016 | https://dplyr.tidyverse.org/ |
| MAST v1.14.0 | Finak 2015 | https://github.com/RGLab/MAST |
| tibble v3.4.0 | Müller et al, 2020 | |
| scVelo v.0.2.2 | Bergen et al, 2020 | https://scvelo.readthedocs.io/index.html |
| ggplot2 v.3.2.1 | Wickham 2016 | Springer-Verlag, https://ggplot2.tidyverse.org |
| Velocyto v0.17 | Manno et al, 2018 | https://velocyto.org/velocyto.py/index.html |
| Tidyverse v1.3.0 | | https://www.tidyverse.org/ |
| FlowJo v10 | | https://www.flowjo.com/ |
| ImageJ v2.1.0 | | https://imagej.net/ImageJ |
| MuscleJ | Mayeuf-Louchart et al, 2018 | |
| TANGO v2.7 | Ollion et al, 2013 | https://imagej.net/3D_ImageJ_Suite |
| **Other** | | |
| Bioinformatics analysis code | This manuscript | https://github.com/larouchej/VML_Dynamics |

### METHODS

**Animals:** C57BL/6 wild-type female and male mice were obtained from Jackson Laboratories or a breeding colony at the University of Michigan (UM). Pax7Cre$^{ER/+}$;Rosa26$^{mTmG/+}$ (mTnG) and Pax7Cre$^{ER/+}$;Rosa26$^{dTomato/+}$ mice were obtained from a breeding colony at UM and administered 5 daily 100uL intraperitoneal injections of 20mg/mL tamoxifen in corn oil. All mice were fed normal chow ad libitum and housed on a 12:12 hour light-dark cycle under UM veterinary staff supervision. All procedures were approved by the University Committee on the Use and Care of Animals at UM and the Institutional Animal Care and Committee and were in accordance with the U.S. National Institute of Health (NIH).

**Injury Model:** Adult mice (3-4 months) were anesthetized with 1.5% isoflurane and administered 0.1mg/kg buprenorphine in 100μL saline. Puralube ointment was applied to both eyes to prevent dryness. Hair was removed from the hindlimbs using clippers, followed by Nair hair removal cream. The surgical area was wiped with Providone Iodine followed by 70% ethanol three times to sterilize. A 1cm incision was made in the skin on the anterior side of each quadricep, followed by the removal of a 2mm or 3mm full depth muscle section from middle of the rectus femoris muscle. In tissues used for histological and immunohistological analysis, a suture was placed in the muscle to identify the middle of the injury. No intramuscular sutures were placed in tissues used for flow cytometry or scRNA-Seq due to the foreign body response induced around the suture. The skin was sutured closed using 7-0 proline sutures, which were removed 7 days post-surgery.

**Histology**

***Tissue Sectioning:*** Quadriceps muscles from both hindlimbs were embedded in optical cutting temperature (OCT) compound and frozen right after excision in isopentane previously cooled with liquid nitrogen then stored at -80˚C. Ten serial cross sections were cut using a cryotome at -20˚C at the suture delineating the midpoint of the injury and collected on positively charged glass slides (Fisher #12-550-15). Duplicate or triplicate cross sections from each timepoint and wound size were placed on the same slide to control for possible staining differences between slides.

***Hematoxylin and Eosin Staining:*** Slides were submerged in hematoxylin for two minutes followed by two sets of ten quick immersions in distilled water. Slides were then submerged in Scott's tap water (20.0g magnesium sulfate heptahydrate and 2.0g sodium bicarbonate in 1L tap water) and distilled water for one minute as well as one set of ten immersions in 80% ethanol. After one minute of Eosin immersion slides were immersed ten times in two different 95% ethanol baths and a 100% ethanol bath. Finally, slides were submerged in SafeClear II for one minute, and two drops of Permount were added before placing the coverslip on top. Brightfield images were taken using a motorized Olympus IX83 microscope at 10X magnification and stitched using the Olympus CellSense software to obtain an image of the complete tissue section.

***Picrosirius Red Staining****:* Slides were removed from -80˚C, thawed to room temperature, and dried for 30 minutes before fixing in 4% paraformaldehyde in PBS for 15 minutes at room temperature. Following fixation, slides were washed 3 times in PBS for 5 minutes each, 2 times in deionized water for 5 minutes each, allowed to dry for 10 minutes at RT, and incubated in 0.5g Direct Red 80 solubilized in 500mL Picric Acid for 1 hour at room temperature. Next, slides were washed twice with acidified water (2.5mL Glacial Acetic Acid in 500mL deionized water) for 5 minutes each followed by 2 washes in deionized water for 5 minutes each. Tissue samples were then dehydrated in a series of ethanol washes (50%, 70%, 70%, 90%, 100%, 100%), incubated in Xylenes twice for 5 minutes each, and mounted with Permount. Brightfield images were taken using a motorized Olympus IX83 microscope at 10X magnification and stitched using the Olympus CellSense software to obtain an image of the complete tissue section. Images were converted to RGB stack and converted to greyscale. To calculate collagen fraction, the green channel was automatically thresholded and measured using ImageJ, then divided by the total muscle cross section area based on thresholding the red channel to include only the tissue.

***Immunohistofluorescence Staining:*** Immunofluorescence staining was performed as previously reported[1]. Briefly, slides were removed from -80˚C, thawed to room temperature for 30 minutes, then fixed in ice-cold 100% acetone at -20˚C for 10 minutes. Following fixation, slides were air-dried for 10 minutes at RT, tissue sections were circled with Hydrophobic Barrier PAP Pen and allowed to dry at room temperature. Tissue sections were the rehydrated in PBS for 5 minutes at RT, then blocked for 1 hour at room temperature using 10% normal goat serum (NGS) in PBS. After blocking, slides were incubated overnight at 4˚C in a solution containing primary antibodies (rabbit anti-laminin (1:500 dilution), goat anti-NKp46/NCR1 (3ug/mL) and/or rat anti-Ly6G (1:500)) diluted in 10% NGS. Primary antibodies were then washed three times for 5 minutes with PBS at room temperature. Secondaries (donkey anti-goat AF488

(1:500 dilution), goat anti-rabbit AF555 (1:500 dilution), goat anti-rat AF647 (1:500), and/or goat anti-mouse AF488 (1:500)) and Hoescht 33342 (1:500 dilution) were added in PBS and incubated for 1 hour at RT. After incubation, slides were washed three times with PBS and a coverslip was mounted using Prolong Diamond florescent mounting medium. Slides were allowed to dry overnight, then stored at 4˚C until imaging. Images were acquired with a Nikon A1 confocal microscope equipped with Colibri 7 solid state light source and pseudo colored using ImageJ. Myofiber area and Feret diameters were quantified based on stitched images of laminin stains using the MuscleJ[2] plugin for ImageJ and R for plotting.

**Extraction and Preparation of Single Cell Suspension from Muscle**

Mouse quadriceps were extracted, and separately weighed using sterile surgical tools and placed into separate petri dishes containing ice-cold PBS. Using surgical scissors, muscle tissues were minced and Collagenase type II (0.2%) and Dispase II (2.5U/mL) were added to 10mL of DMEM per quadricep. Samples were placed on rocker in a 37˚C incubator for 1 hour and mixed by pipette every 30 minutes. The enzymes within the slurry were then inactivated by addition of 20% heat-inactivated fetal bovine serum (HI-FBS) in Ham's F10 media. The solution was passed through a 70um cell strainers, centrifuged, washed, and counted.

**Flow Cytometry**

The single cell suspension was pelleted and re-suspended in staining buffer (PBS with 2% BSA, 2mM EDTA and 0.01% sodium azide) at 2E8 cells/mL, then 200uL of cells were plated into round-bottom 96-well plates for staining. Cells were centrifuged at 250xg for 2.5 minutes. The supernatant was discarded and cells were re-suspended in a primary antibody cocktail that included one or more of the following: CD11b-PE (1:200 dilution), CD11b-APC (1:200 dilution), CD45-AF488 (1:100 dilution), CD68-FITC (1:400 dilution), APC-CD335 (NKp46) (1:40 dilution), Ly-6A/E-APC (1:200), CD45-APC (1:200), Ter119-APC (1:200), CD29-PE (1:100), CD184-biotin (1:50), and/or BV421-Ly6G (1:400 dilution). Cells were incubated in primary antibody for 30 minutes on ice, then centrifuged, washed with staining buffer, and re-suspended staining buffer containing PECy7 streptavidin (1:100 dilution) where applicable. Cells were incubated with the secondary antibody for 20 minutes on ice, washed with staining buffer, then re-suspended in staining buffer containing propidium iodide (1:400 dilution) for 1 minute at room temperature in the dark (where appropriate), centrifuged, and re-suspended in staining buffer for flow cytometry analysis. Prior to acquisition, cells were filtered through 40μm cell strainers to prevent cytometer clogging. Single color controls were made using UltraComp eBeads compensation beads stained according to manufacturer's protocol. Florescent minus one controls were made using a mix of cells from each sample. For analysis of apoptosis, cells were stained with Annexin V immediately prior to flow cytometry analysis according to manufacturer's instructions. Samples were acquired within 1 hour on a BioRad Ze5 cytometer, and the data was processed using FlowJo (version 10) with manual compensation.

**Real-Time PCR (qRT-PCR)**

Cell lysates were thawed at room temperature for 30 minutes, then RNA was extracted using the Qiagen miRNeasy Micro Kit according to manufacturer's instructions. RNA purity and concentration were measured using a NanoDrop and Qubit RNA HS Assay. Within one week, cDNAs were synthesized using the SuperScript III cDNA Synthesis Kit according to manufacturer's protocol. DNA quality and concentration was determined using a NanoDrop. 80-100ug cDNA template was plated in triplicate along with SYBR Green PCR MasterMix and 500nM PCR primer, then cycled 40 times starting at 95˚C for 10 seconds followed by 60˚C for 30 seconds on a CFX96 Real-Time thermocycler. Gene expression was quantified using the $\Delta\Delta$Ct method.

**NK Cell and Neutrophil Co-Culture**

Murine IL15 super agonist (mIL15SA) was complexed as previously described[5], aliquoted in sterile PBS with 0.1% BSA, and stored at -80˚C until use. Splenic NK cells were magnetically enriched from uninjured young WT mice using Miltenyi mouse NK Cell Isolation Kit per manufacturer's instructions and cultured for 48 hours in RPMI containing 10% FBS, 60ng/mL mIL15SA, and antibiotics to activate. NKp46$^{+}$ NK cell purity following magnetic enrichment was determined to be >80% by flow cytometry (Supp. Fig. 4b). Neutrophils were FACS-enriched (CD11b+Ly6G+PI-) from mouse skeletal muscle 7 days post 3mm VML injuries. NK cells and neutrophils were co-cultured at a 10:1 ratio for 4 hours at 37˚C in RPMI with 10% FBS and antibiotics. Following incubation, cells were stained with Annexin V and PI according to manufacturer's protocol, then immediately analyzed on a BioRad Ze5 flow cytometer. Data was analyzed using FlowJo.

**NK Cell Transplants**

Splenic NK cells were obtained as described above and activated in vitro using IL15SA for 48 hours. Activated NK cells were washed and re-suspended in sterile PBS (150K cells/10μL) and transplanted into 2mm VML defects 7 days post injury using a 29 ½ G needle. Contralateral limbs were used as controls and received PBS only. At 14dpi, mice were euthanized, and neutrophil abundance was quantified by flow cytometry ($CD11b^+Ly6G^+PI^-$). At 28-dpi, a separate cohort of mice were humanely euthanized, and healing was analyzed histologically to quantify % collagen, muscle mass, and myofiber diameter as described above.

**CCR1 Inhibition**

Mice were administered bilateral 3mm VML defects. Starting 7dpi mice received daily IP injections of CCR1 inhibitor J-113863 (8mg/kg) solubilized in sterile PBS with 5% DMSO and 5% CremophorEL. Neutrophil abundance was quantified by flow cytometry ($PI^-CD11b^+Ly6G^+$) at 14-dpi.

**TGFβRII Inhibition**

Mice were administered bilateral 3mm VML defects to the quadriceps, or 2mm VML defects to the tibialis anterior for force testing. Starting 3-dpi mice received intramuscular injections of ITD1 (200pg in 20μL using a 29 ½ G needle) in sterile PBS with 0.01% DMSO every third day until 15-dpi. Contralateral limbs were injected with 20uL of vehicle (PBS with 0.01% DMSO) every third day. Neutrophil abundance was quantified by flow cytometry at 14-dpi, and healing was assessed histologically at 28-dpi as described above.

**Muscle Force Testing**

Force testing was performed as previously reported by Dellorusso et al. with some modifications[6]. Mice were anesthetized with 2% Isoflurane to maintain a deep anesthesia throughout the experiment. Hindlimb fur was removed with animal clippers. The TA muscle was exposed by removing the overlying skin and outer fasciae. The distal TA tendon was isolated, and the distal half of the TA was freed from adjacent muscles by carefully cutting fasciae without damaging muscle fibers. A 4–0 silk suture was tied around the distal tendon, and the tendon was severed. The animal was then placed on a temperature-controlled platform warmed to maintain body temperature at 37°C. A 25-gauge needle was driven through the knee and immobilized to prevent the knee from moving. The tendon was tied securely to the lever arm of a servomotor via the suture ends (6650LR, Cambridge Technology). A continual drip of saline warmed to 37°C was administered to the TA muscle to maintain its temperature. The TA muscle was initially stimulated with 0.2 ms pulses via the peroneal nerve using platinum electrodes. Stimulation voltage and muscle length were adjusted for maximum isometric twitch force (Pt). While held at optimal muscle length (Lo), the muscle was stimulated at increasing frequencies until a maximum force (Po) was reached, typically at 200 Hz, with a one-minute rest period between each tetanic contraction. Subsequently, the same procedure was repeated, but rather than activating the muscle via the peroneal nerve, a cuff electrode was placed around the muscle for stimulation. Muscle length was measured with calipers, based on well-defined anatomical landmarks near the knee and the ankle. Optimum fiber length (Lf) was determined by multiplying Lo by the TA Lf/Lo ratio of 0.6[7].

After the evaluation of isometric force, the TA muscle was removed from the mouse. The tendon and suture were trimmed from the muscle, and the muscle was weighed. After removal of TA muscles, deeply anesthetized mice were euthanized by the induction of a pneumothorax. Total muscle fiber cross-sectional area (CSA) of TA muscles was calculated by dividing muscle mass by the product of Lf and 1.06 $mg/mm^3$, the density of mammalian skeletal muscle[8]. Specific Po was calculated by dividing Po by CSA.

**Muscle Progenitor Cell Differentiation**

***Primary Cells:*** After obtaining a single cell suspension, magnetic enrichment of MuSCs was performed using the mouse Satellite Cell Isolation Kit according to manufacturer's instructions. Meanwhile, a 10cm tissue culture dish was coated with 10% Matrigel in DMEM as previously published[9]. Enriched MuSCs were seeded onto the dish in 8mL myoblast medium (Ham's F10 containing 20% HI-FBS, 10ug/mL fibroblast growth factor (bFGF), and 1X penicillin/streptomycin). Cells were expanded for 5 days with fresh media added every other day. Following expansion, 50,000K cells were passaged using 0.25% trypsin and seeded into each well of a Matrigel-coated 12-well plate and expanded until 90-100% confluency in myoblast media. At this point, myoblast media was replaced with differentiation media (DMEM with 5% horse serum and 1% penicillin/streptomycin) and allowed to differentiate for 72 hours.

***Immunofluorescence Staining:*** After 72 hours in differentiation media, myotubes were rinsed twice with PBS, then fixed with 4% paraformaldehyde for 20 minutes at room temperature (RT). After three quick washes with 1X PBS, cells were incubated for 15 minutes at RT with 0.1% TritonX-100, washed three times with 1X PBS, and blocked for 1 hour at RT with 1% BSA, 0.1% Tween-20 and 22.52mg/mL glycine in 1X PBS. After blocking, cells were incubated with primary antibodies (1:10 dilution of anti-MYH3) overnight at 4˚C followed by secondary antibodies (1:500 dilution of AF555 anti-rabbit) overnight at 4˚C. Following secondary stain, cells were washed three times in 1X PBS, then nuclei were stained with Hoechst 33342 (1:500 dilution). Immunolabeled cells were imaged on a Zeiss epifluorescent microscope using a 20X objective. Ten images were acquired and analyzed per well. The fusion index was automatically calculated using MATLAB as the ratio of nuclei within myofibers containing more than 2 nuclei divided by the total number of nuclei per image. Myonuclei as a percentage of total cells were calculated by hand as the number of MyH3+ myotubes with more than 2 nuclei divided by the total number of cells.

***Real-Time PCR:*** After 72 hours in differentiation media, myotubes were rinsed twice with PBS then collected into Qiazol, vortexed for 1 minute, incubated for 5 minutes at room temperature, and stored at -80˚ until further analysis. qRT-PCR was performed as described above.

**Single-cell mRNA Sequencing**

***Sample Preparation and Sequencing:*** Freshly isolated cells were briefly stained with Propidium Iodide (PI) and FACS sorted to remove dead cells (PI positive) and debris. An equal number of viable cells were pooled from two mice and 10,000-16,000 cells were loaded into the 10x Genomics chromium single cell controller and single cells were captured into nanoliter-scale gel bead-in-emulsions (GEMs). cDNAs were prepared using the single cell 3′ Protocol as per manufacturer's instructions and sequenced on a NovaSeq 6000 (Illumina) with 26 bases for read1 and 98x8 bases for read2. With the exception of 7dpi, which was repeated twice pooling two mice of the same gender, sequencing libraries were generated from a pool of cells from one male and one female mouse.

***Data Processing and Analysis:*** 10x CellRanger v2.0.0 software's mkfastq and count command were run with default parameters except expect-cells=10000. HDF5 matrix files were imported into R (https://www.r-project.org/) using the Seurat v3 package[10] and genes expressed in less than 200 cells or cells expressing less than 3 genes were removed. Seurat objects were merged and cells with >10% mitochondrial genes or over 7500 reads were deemed low quality and removed. Following normalization, identification of variable features, and scaling using default parameters, datasets were integrated using Seurat's FindIntegrationAnchors (dims = 1:30, k.filter = 200, k.score = 30, anchor.features = 2000) and IntegrateData functions. Dimensional reduction was performed in Seurat using PCA, then UMAP[11] followed by community detection using the Louvain algorithm (resolution = 0.5). Cluster marker genes were determined both with Seurat (only.pos = T, min.pct = 0.25, logfc.threshold = 0.25, return.thresh = 0.01) and scCATCH (species = 'Mouse', match_CellMatch = T, tissue = 'Muscle', cell_min_pct = 0.25, logfc = 0.25, pvalue = 0.05) to annotate cell types. Defect-associated differential gene expression among celltypes was calculated using MAST[12] as previously reported[13]. NicheNet[14] interaction predictions were generated using the Seurat Wrapper according to the vignette to compare the 3mm (degenerative) defects to the 2mm (regenerative) defects. Seurat and Ggplot2 were used for data visualization. Gene expression and cell type abundances were similar across libraries generated from male and female mice 7dpi (data not shown).

**Statistics**

Experiments were repeated at least twice, with the exception of scRNA-Seq. Bar graphs show mean ± standard error unless otherwise stated. Statistical analysis was performed in MATLAB and/or R using two-sample Student's t-test assuming normal distribution and equal variances, one-way ANOVA, paired-T-test, or Kolmogorov-Smirnov test for distributions, as specified in figure captions. All statistical tests performed were two-sided. Outliers were determined using the Tukey's fences method with k = 1.5 and removed from further analysis. P-values less than 0.05 were considered statistically significant. Effect sizes were estimated using Cohen's d.
